# Allosteric perspective on the mutability and druggability of the SARS-CoV-2 Spike protein

**DOI:** 10.1101/2021.08.01.454696

**Authors:** Zhen Wah Tan, Wei-Ven Tee, Firdaus Samsudin, Enrico Guarnera, Peter J. Bond, Igor N. Berezovsky

**Affiliations:** Bioinformatics Institute, Agency for Science, Technology and Research (A*STAR), 30 Biopolis Street, #07-01, Matrix, 138671, Singapore; Department of Biological Sciences (DBS), National University of Singapore (NUS), 8 Medical Drive, 117579, Singapore

**Author notes:** Global Analytical Pharmaceutical Science and Innovation, Merck KGaA, Via Luigi Einaudi, 11, Guidonia Montecelio - 00012 Rome, Italy (Enrico Guarnera). These authors contributed equally to the paper (Zhen Wah Tan, Wei-Ven Tee). To whom correspondence should be addressed: Igor N. Berezovsky.

## Abstract

Recent developments in the SARS-CoV-2 pandemic point to its inevitable transformation into an endemic disease, urging both diagnostics of emerging variants of concern (VOCs) and design of the variant-specific drugs in addition to vaccine adjustments. Exploring the structure and dynamics of the SARS-CoV-2 Spike protein, we argue that the high mutability characteristic of RNA viruses coupled with the remarkable flexibility and dynamics of viral proteins result in a substantial involvement of allosteric mechanisms. While allosteric effects of mutations should be considered in predictions and diagnostics of new VOCs, allosteric drugs advantageously avoid escaping mutations via non-competitive inhibition originating from many alternative distal locations. The exhaustive allosteric signalling and probing maps provide a comprehensive picture of allostery in the Spike protein, making it possible to locate sites of potential mutations that could work as new VOCs “drivers”, and to determine binding patches that may be targeted by newly developed allosteric drugs.

## Introduction

Allostery is a universal property of all proteins and protein machines regardless of their structures, sizes, or functions (Gunasekaran et al., 2004; Mitternacht and Berezovsky, 2011), by which proteins recognize environmental cues in the form of perturbations (Guarnera and Berezovsky, 2019a), such as binding of small ligands (Guarnera and Berezovsky, 2016a), mutations (Guarnera and Berezovsky, 2020; Tee et al., 2019), post-translational modifications (Berezovsky et al., 2017; Mitternacht and Berezovsky, 2011), and elicit a response at remote locations (Guarnera and Berezovsky, 2019a; Tee et al., 2021). The ectodomain of the spike (S) glycoprotein is a large interlocking molecular machine that undergoes extensive conformational rearrangements in order to fuse the viral and cell membranes (Walls et al., 2020) upon attachment of the receptor-binding domain (RBD) to the host receptor (Wrapp et al., 2020). The orchestration and the regulation of these complex processes are facilitated by the S protein modular (Gobeil et al., 2021; Wrapp et al., 2020) and hinged (Turonova et al., 2020) structure, conformational dynamics of which are regulated through communication between distant subunits, hinting thus at the existence of allosteric communication and signalling between them (Gobeil et al., 2021; Raghuvamsi et al., 2021).

There are several sequence/structure determinants of the SARS-CoV-2 S protein dynamics an interactions that might be linked to the virus infectivity and transmissibility. For example, while having 98% of sequence identity with the S protein from the closely-related bat coronavirus RaTG13, it has 29 variant residues with 17 of them located in the RBD (Walls et al., 2020; Wrapp et al., 2020; Wrobel et al., 2020), which might be one of the reasons for the 10-20 fold higher affinity in its binding to the human angiotensin-converting enzyme 2 (ACE2) than that of SARS-CoV S protein (Wrapp et al., 2020). The SARS-CoV-2 S protein also reveals higher thermostability (Wrobel et al., 2020) and possesses a polybasic “RRAR” insertion (instead of a single “R” in the corresponding site of SARS-CoV S), resulting in an S1-S2 furin cleavage site potentially flanked by three additional O-linked glycans unique for SARS-CoV-2 (Andersen et al., 2020). The site is associated with increased pathogenicity of SARS-CoV-2 (Wrobel et al., 2020), of related coronaviruses, including HKU1 (Chan et al., 2008) and MERS-CoV (Millet and Whittaker, 2014, 2015), and of other viruses such as influenza virus (Steinhauer, 1999). In total there are 22 N-linked glycosylation locations per protomer (Watanabe et al., 2020a; Watanabe et al., 2020b); 9 of the 13 glycans in S1 and all 9 glycans in S2 are conserved among SARS-CoV-2 and SARS-CoV (Walls et al., 2020).

Multiple conformational transitions between states with potentially different epitopes including the immunodominant, non-neutralizing ones, further complicated by the possibility of an antigenic drift in the eventually endemic SARS-CoV-2, present challenges for the vaccine design and development of therapeutic approaches against this infectious disease (Cai et al., 2020). The urgent necessity to expand therapeutic options for COVID-19, in addition to vaccines, is further accentuated by the findings from a large randomized control trial coordinated by the World Health Organization, which indicated that the repurposed drugs remdesivir, lopinavir/ritonavir, hydroxychloroquine and interferon β-1a confer little to no anti-viral effects on hospitalized patients (Pan et al., 2020). Starting from traditional approaches, such as antibodies and their cocktails (Chi et al., 2020; Li et al., 2020a), the quest for therapies has navigated towards the use of convalescent sera (Zhou et al., 2020), nanobodies (Huo et al., 2020), and to the de novo design of miniprotein inhibitors (Cao et al., 2020). The major obstacle in both vaccine and therapeutics developments (Gobeil et al., 2021) is the constantly evolving, with characteristically high level of mutations and continuous evolution of the RNA virus (Harvey et al., 2021). While much attention has been placed on the mutations in the RBD, there are ever increasing data showing that mutations may work indirectly, affecting remote units of the S protein. A notable example is the D614G mutation of the S protein, which became dominant concomitant with its emergence (Korber et al., 2020). Although its implications of this mutation are still under debate (Grubaugh et al., 2020), a study by Yurkovetskiy *et al*. has shown that the mutation, which is located outside the receptor-binding domain (RBD), causes the S protein to favor an open conformation (Yurkovetskiy et al., 2020). The “mutability” challenge motivated a high-throughput analysis of the effects of mutations (Li et al., 2020a) with the goal “to predict their effect rather than just to seeing it” (Starr et al., 2021; Starr et al., 2020). Since a number of mutations apparently act indirectly, implicating an allosteric mode of action (Gobeil et al., 2021; Li et al., 2020a; Starr et al., 2021; Starr et al., 2020), our objective here is to quantify the energetics of the allosteric modulation caused by these mutations, to determine patterns of relevant signalling and locations of potential druggable sites for therapeutic interventions.

In this work we perform a comprehensive analysis of allosteric signalling that may be involved in the regulation of the S protein activity and its modification upon rapid viral evolution (Harvey et al., 2021). Contrary to previous works discussing circumstantially detected cases of allosteric communication (Gobeil et al., 2021; Raghuvamsi et al., 2021) caused by certain mutation(s), we use our computational framework that allows to perform a comprehensive analysis of allosteric signalling on the basis of the structure-based statistical mechanical model of allostery (SBSMMA). The SBSMMA (Guarnera and Berezovsky, 2016b) is an exact model providing the per-residue free energy of allosteric signalling upon ligand/probe binding, mutations and their combinations, allowing to build exhaustive allosteric signalling maps (ASMs, (Guarnera and Berezovsky, 2019b)) and allosteric probing maps (APMs, (Tan et al., 2020)). On the basis of the extensivity of the free energy, we show how high-throughput data from ASMs/APMs data can be used for predicting allosterically acting mutations that may lead to new VOCs, as well as for finding targets for allosteric drug development. We discuss a generic strategy for the analysis of the allosteric effects in future diagnostics of new variants, in avoiding the effects of escaping mutations, in design of new medicines, and in preventing the emergence of drug resistance in the “cocktail” therapeutics (Chi et al., 2020).

## Results

### Allosteric signalling in the pre-fusion S glycoprotein upon simulated receptor binding

The S glycoprotein from SARS-CoV-2 is a large complex of three interlocking monomers that plays a critical role in viral entry into the host cell by binding to the human ACE2 receptor (Walls et al., 2020; Yan et al., 2020). It consists of the S1 and S2 subunits (Figure 1A), which are involved in the recognition and binding to the host receptors, and the subsequent membrane fusion, respectively (Walls et al., 2020; Wrapp et al., 2020). Binding to host receptors triggers large structural rearrangements in the metastable pre-fusion conformation of the S protein, resulting in the shedding of S1 and, in turn, fusion of the viral envelope with the cell membrane mediated by the S2 core(Li, 2016). Figure 1A shows the S glycoprotein (residues 27-1213) in the open state, in which the receptor-binding domain (RBD) of one of the monomers (chain A, orange) is pointed upward to engage a receptor, hereinafter referred to as the “up” conformation. The RBD of chains B and C (colored in green and purple) adopt the “down” conformation inaccessible to receptors. In the closed state, all RBDs adopt the down conformation, capping the top of the S2 subunits (Figure S1). The S1 subunit consists of the RBD and the N-terminal domain (NTD), which are respectively associated with the subdomains 1 and 2 (SD1 & SD2) (Figure 1A). The SD1 and SD2 together provide an interface between the S1 subunit and the S2 subunit of another monomer. The predominantly alpha-helical S2 core (Figure 1A) comprises various regions including the fusion peptide (FP), which is inserted into the host cell membrane after structural rearrangements triggered by receptor binding(Bosch et al., 2003). In this study, we modelled the ectodomain of the pre-fusion S glycoprotein in the open and closed states based primarily on available structures (Walls et al., 2020; Wrapp et al., 2020) obtained from cryo-electron microscopy (cryo-EM). The SBSMMA (Guarnera and Berezovsky, 2016b) allows to determine individual residues and sites that undergo positive or negative allosteric modulation as a result of a perturbation, such as mutation(s), binding, glycosylation, or different combinations thereof. The positive allosteric modulation (blue) points to potential conformational changes as a result of the work exerted on the residue/site, whereas the negative allostery (red) reveals stabilization of modulated residues/sites that dampens their dynamics.

**Figure 1.**
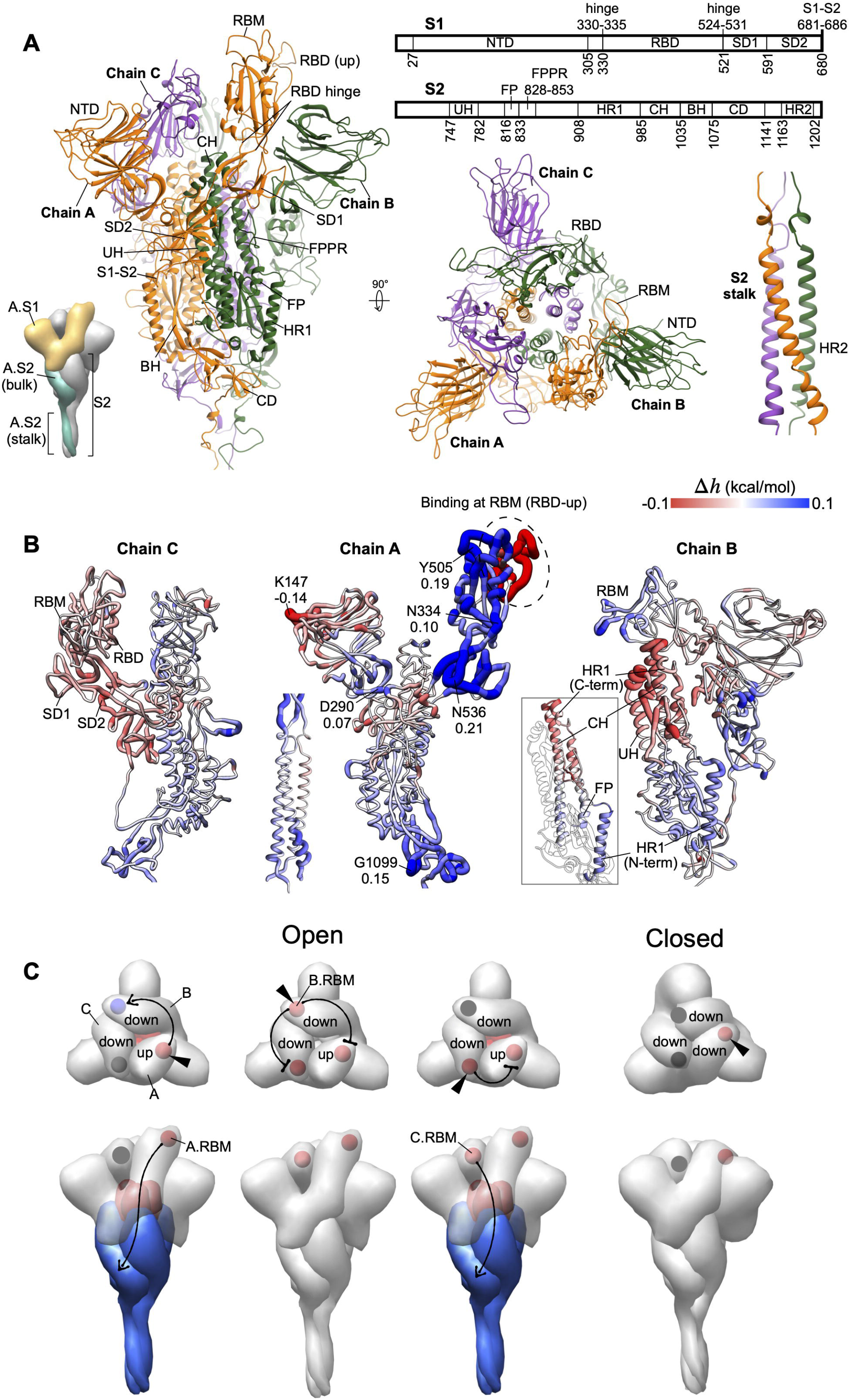
Allosteric response upon simulated receptor binding. **(A)** The structure of the spike homotrimer in the open state modelled based on a cryo-EM structure (PDB: 6VSB). Chains A-C are colored in orange, green and purple, respectively. The residues forming each region are defined based on other work (Cai et al., 2020; Wrapp et al., 2020). NTD: N-terminal domain, RBD: receptor-binding domain, SD1: subdomain 1, SD2: subdomain 2, RBD hinge: two loops linking RBD and SD1, S1-S2: S1-S2 cleavage site, UH: upstream helix, FP: fusion peptide, FPPR: fusion peptide proximal region, HR1: heptad repeat 1, CH: central helix, BH: beta hairpin, CD: connector domain, HR2: heptad repeat 2. **(B)** The allosteric modulation (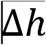, kcal/mol) at every residue due to the binding at A.RBM (circled) is depicted using the color-gradient presentation from positive (conformational changes, blue) to negative (stabilization, red) modulation. The complete data on binding to each RBM in the open or closed state is shown in Figures S1B and S1C. **(C)** The schematic shows the allosteric communication between RBMs (spheres) of different monomers and from the RBM to the S2 subunit upon ACE2 binding. In each scenario, the bound RBM is indicated by a triangle symbol, and the sign of the modulation in the RBM and S2 is colored in blue, red or grey for positive, negative, or very weak/absent modulation, respectively.

To explore the allosteric responses originated by the interactions/binding with/to a host receptor, we simulated the binding at the receptor-binding motif (RBM, defined in Figure S1A) in the RBD. Figure 1B shows the allosteric modulation of the open S homotrimer upon binding of chain A’s RBM (A.RBM, circled) in the RBD-up conformation. Binding to the A.RBM results in a positive allosteric modulation, originating a configurational work exerted in several regions of chain A, indicating, thus, their potential conformational changes (Figure 1B): the RBD core, SD1, the loops joining the RBD and SD1 (referred to as RBD hinge), the region linking NTD with SD2, and the lower part of S2, especially in the connector domain (CD). The residues under the strongest allosteric modulation in each of these regions are indicated with the modulation values (Figure 1B, middle structure). This observation is consistent with a cryo-EM study, which revealed the rigid-body rotation of the up RBD away from the trimer axis and the shift of the centers of mass of NTD, SD1 and SD2 upon binding of S protein to ACE2 (Benton et al., 2020). The large and uniform increase in the configurational work exerted predominantly in the RBD and SD1 compared to elsewhere is further corroborated by the dramatic opening of the spike observed in the exascale molecular dynamics simulations (Zimmerman et al., 2021), which is only possible through substantial movements of these two domains.

Shifting focus to modulation occuring in chain B as a result of the binding at A.RBM, we observed a configurational work exerted at B.RBM, likely causing conformational changes in the B.RBD (Figure 1B, right structure), thus yielding allosteric communication between the distant RBMs of chains A and B (Figure 1B). Noteworthy, the S2 subunit of chain B, packed next to A.SD1 and A.SD2, displays opposite allosteric modulations in its upper and lower portions. The N-terminal end of the heptad repeat 1 (HR1) and FP in the lower portion are positively modulated, whereas the C-terminal end of HR1 and parts of the central helix (CH) and upstream helix (UH) located in the upper portion are negatively modulated preventing its conformational changes (Figure 1B, inset). The observation that binding at the A.RBM produces a positive configurational work exerted at the N-terminal end of B.HR1 and a concomitant negative modulation at the other end of B.HR1 (Figure 1B, inset) is in a good agreement with previous studies on the S protein dynamics (Walls et al., 2017). Specifically, it was shown that the HR1 undergoes a jackknife-like movement in the transition from the pre-fusion to the post-fusion state, in which the N-terminal part of HR1 swings upward and reorients to form a continuous α-helix with the adjoining central helix (Cai et al., 2020; Walls et al., 2017). Moreover, the fusion peptide (FP), a critical element in the fusion machinery, which is also known to undergo large structural rearrangements, as well as the S2’ cleavage site located immediately upstream, yields positive modulation (Figure 1B, right), albeit slightly weaker compared to the other adjacent regions. The C.RBD (excluding the C.RBM) of chain C, C.SD1, and C.SD2 show negative allosteric modulation, and the C.S2 subunit is chiefly positively modulated (Figure 1B, left structure). The S2 coiled coil stalk formed by the heptad repeat 2 (HR2) region of all three monomers is positively modulated at both ends of the stalk, *i*.*e*. next to the CD and near the transmembrane domain.

Summarizing the above observations, one can conclude that binding at the A.RBM induces an allosteric response in several locations of both S1 and S2 subunits in all chains of the S protein, amongst which the RBD and the adjacent SD1 in S1 exhibit the largest increase in configurational work (Figure 1B), which is likely to result in conformational changes that lead to the dissociation of the monomeric S1 domain upon binding (Benton et al., 2020). While the down conformation of the RBD is commonly deemed inaccessible for receptor binding, from a practical viewpoint, some neutralizing antibodies bind to the RBD in the down as well as the up conformations (Barnes et al., 2020), calling thus for an investigation of the allosteric signalling emanated from the perturbed down RBDs with potential implications for antiviral antibody design (Samsudin et al., 2020). Therefore, in order to obtain a complete picture of the allosteric communication between the RBMs and other function-related locations we simulated binding to the RBMs of both up and down RBDs in the open (1up-2down) and closed (3down) states (Figure S1 B, C). The modulation modes of allosteric signalling to the other RBMs and the S2 subunit in different scenarios are summarized in Figure 1C. In the open state, binding to A.RBM in the up A.RBD causes positive modulation in B.RBM and the lower part of S2, and negative modulation in the upper part of S2, whereas C.RBM is unaffected, as described above (Figure 1B). Perturbing B.RBM in the down B.RBD leads only to negative modulation in A.RBM and C.RBM. Binding at C.RBM of down C.RBD stabilizes the A.RBM, and, interestingly, leads to extensive allosteric modulation in the S2 subunit (Figures 1C, S1B), similar to that caused by the bound A.RBM. Despite both B.RBD and C.RBD adopting the down conformation, the differences in the signalling from their binding motifs could be attributed to the arrangement of the monomers in the open S structure: in this case the binding motif in chain B is oriented towards the down C.RBD, whereas the binding motif in chain C faces the up A.RBD. This suggests that the allosteric signalling originated from a bound RBM can be affected by the conformation (up/down) and interaction with the RBD in other monomers. In the closed state, in which all three RBDs adopt the down conformation, simulated binding to the RBM of each of the monomers is generally unable to induce allosteric responses in the rest of the S homotrimer (Figure 1C, right structure; Figure S1C).

### Allosteric communication between important structural units of the spike glycoprotein

In order to obtain details of allosteric signalling, we derived the allosteric signalling maps (ASMs, (Guarnera and Berezovsky, 2019b)) of the open and closed states of the spike glycoprotein, which provide exhaustive pictures of allosteric signalling in the form of an allosteric modulation range from every protein residue to the rest of the structure regardless of the type of the original amino acid (see also STAR* Methods for the formal definition of the allosteric modulation range and for the description of the ASM). In this work, we use ASMs with the allosteric modulation range obtained by modelling each residue’s perturbation in the form of replacement from the smallest (Ala/Gly-like) to the bulkiest (Trp/Phe-like) amino acids, mimicking the residue stabilization as a result of a mutation or of an effector binding (see also STAR* Methods for further explanations). The ASMs for both open and closed S structures and the pairwise distance map are shown in Figure S2. The complete, downloadable ASM data is accessible through the AlloMAPS database (Tan et al., 2019) (link to the complete data will be available upon peer-review publication) with the data available for download. In order to delineate the allosteric communication in this large oligomer, we used a top-down approach by first identifying the intra- and inter-monomer communication between protein sites/regions (defined in Figures 1A, S1A), followed by examining the communication at the single-residue resolution. In order to explore the signalling between different chains of the S oligomer we use here sequential ASMs where mutations of all residues are made sequentially in every chain (see STAR* Methods for further explanations).To gain a picture of the allosteric communication between regions, we averaged the modulation range of signalling originated from every residue in a region to each residue in another region and obtained the matrices of allosteric signalling between the S protein regions for the open and closed states (Figures 2A, S2B).

**Figure 2.**
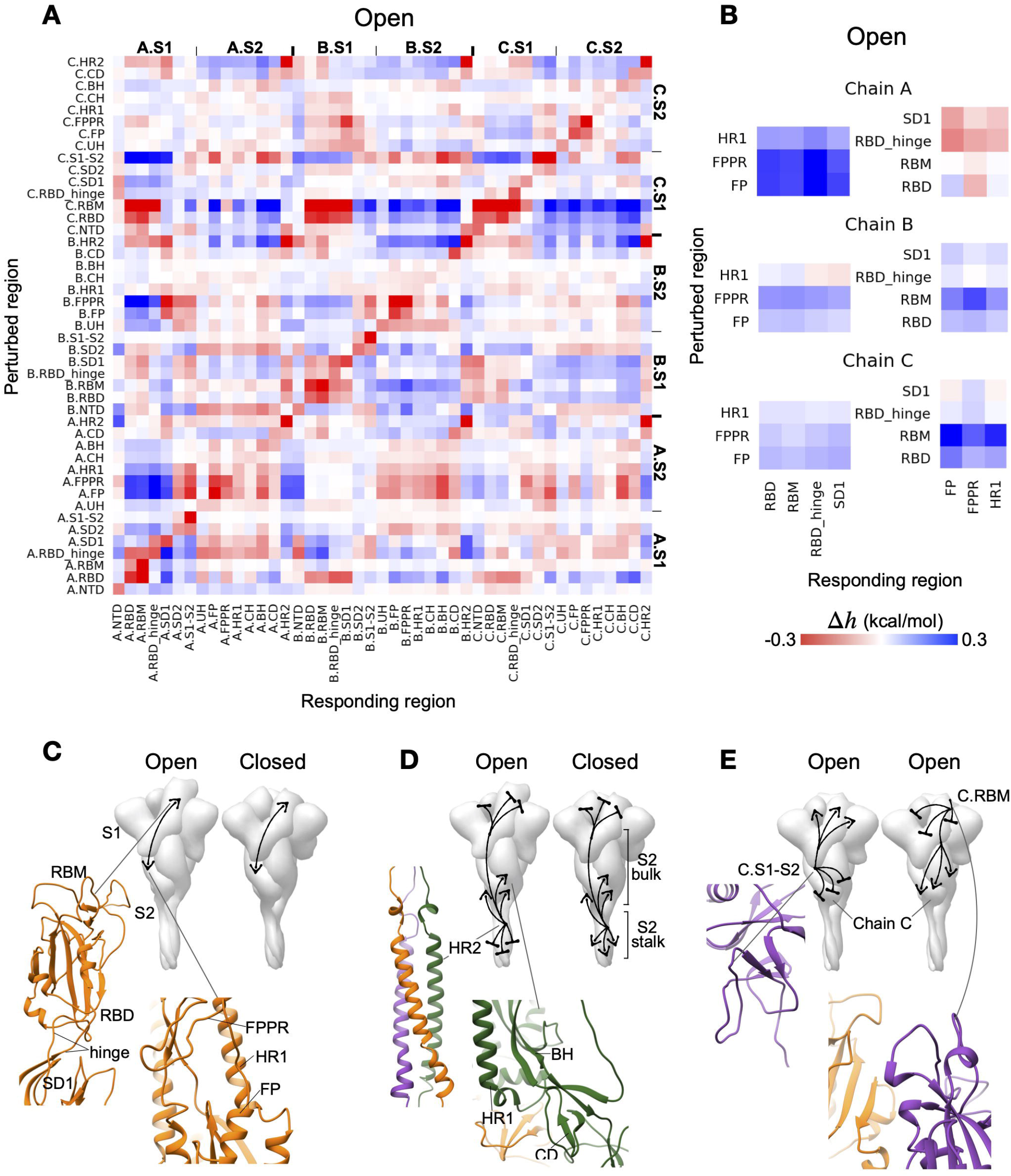
Allosteric communication between structural units. **(A)** ASM containing the average modulation ranges (in kcal/mol) between S protein regions in the open state (the ASM for the closed S is shown in Figure S2B). **(B)** Parts of the complete ASM in (A) highlighting the intra-monomer communication in chains A-C between two clusters of regions: (i) FP/FPPR/HR1 and (ii) RBD/RBM/RBD hinge/SD1. **(C)** Allosteric communication between the two clusters. (D and E) Example of intra- and intermonomer signalling observed in (A). The corresponding parts of the ASM are shown in Figure S2C and S2D.

Figure 2B highlights the signalling between the perturbed FP, the fusion peptide proximal region (FPPR, (Cai et al., 2020)), and HR1 in the S2 subunit and the responding RBD, RBM, RBD hinge, and SD1 units in the S1 subunit of the same monomer in the open S structure. The intra-monomer allosteric modulation linking residues in these two sets of regions in S1 and S2 are predominantly positive in both open and closed states (Figures 2B, 2C and S2B). While the mode and the magnitude of modulations in chains B and C are similar to each other, perturbations in FP, FPPR and HR1 in chain A produce a positive configurational work (Figure 2B, top left) and, potentially, conformational changes in RBD, RBM, RBD hinge, and SD1, and conversely, perturbations in the latter set of regions lead to prevention of structural changes in the FP, FPPR, and HR1 (Figure 2B, top right). In another example, perturbations in the HR2 of a monomer generally result in negative modulation in RBD, RBM, RBD hinge and SD1 in all monomers of both open/closed forms (here and below marked by bases in illustrations of the S-structure), but positive modulation (marked by arrows) in most of the regions in the S2 subunit such as HR1, BH and CD (Figures 2A, 2D and S2C). A notable difference between the signalling patterns observed in the open and closed states is in the modulation of the long S2 stalk (residues 1163-1202, ∼60 Å long): the HR2 regions forming the coiled coil are negatively and positively modulated in the open and closed states, respectively (Figure S2C). In addition to the patterns of signalling that are largely similar across different monomers and conformations (open/closed), we also observed allosteric signals specific only to a certain monomer(s) and conformation. For instance, in the open state, allosteric signalling originated from the S1-S2 cleavage site of chain C (C.S1-S2) promotes conformational changes in all C.S1 regions except C.SD2, while simultaneously precluding conformational changes in the entire C.S2 subunit (Figures 2A, 2E (left structure), S2D). Stabilization of the C.RBM, on the other hand, causes negative modulation in RBD, RBM, RBD hinge and SD1 and positive modulation in S2 in all monomers (Figures 2E (left structure), S2D). In these two cases, the allosteric responses are observed in regions of chain C induced by signals emanating from the S1-S2 and RBM in the chain C of the open spike structure, whereas perturbations of these regions in other chains do not result in the same pattern of signalling in them.

### Allosteric signalling to the receptor-binding motif of chain A (A.RBM)

The comprehensive description of the signalling in the ASMs allows us to explore the locations that may elicit a desired modulation at a site/region of functional importance, hence providing a starting point for allosterically-regulated targeting of functional sites. Since chain A undergoes the most significant conformational changes, we focused here on the allosteric communication to A.RBM from distant regions in the open (Figure 3A) and closed (Figure 3C) conformations (see also Figure S3 for the complete data on signalling to A.RBM). Figure 3 shows that the RBM undergoes different modes of modulation, depending on the location of the perturbation, and adjacent sites/regions usually induce comparable modulations in the RBM. For instance, perturbations in the closely located C.NTD and B.SD1 cause negative modulation in the A.RBM (Figure 3A). Likewise, perturbing A.FP, A.FPPR, A.HR1 and C.SD1 results in a positive modulation in A.RBM. The modulation of A.RBM is summarized in Figure 3 (center) by mapping the allosteric signals from various regions of the spike ectodomain upon single mutations. Comparison of the responses in the open and closed structures reveals common features — negative modulation (marked by bases) from locations in S1 including C.NTD, C.RBM and other neighboring regions, and HR2 in the stalk; positive modulation from A.FP and C.S1-S2 (marked by triangles in Figures 3A, C).

**Figure 3.**
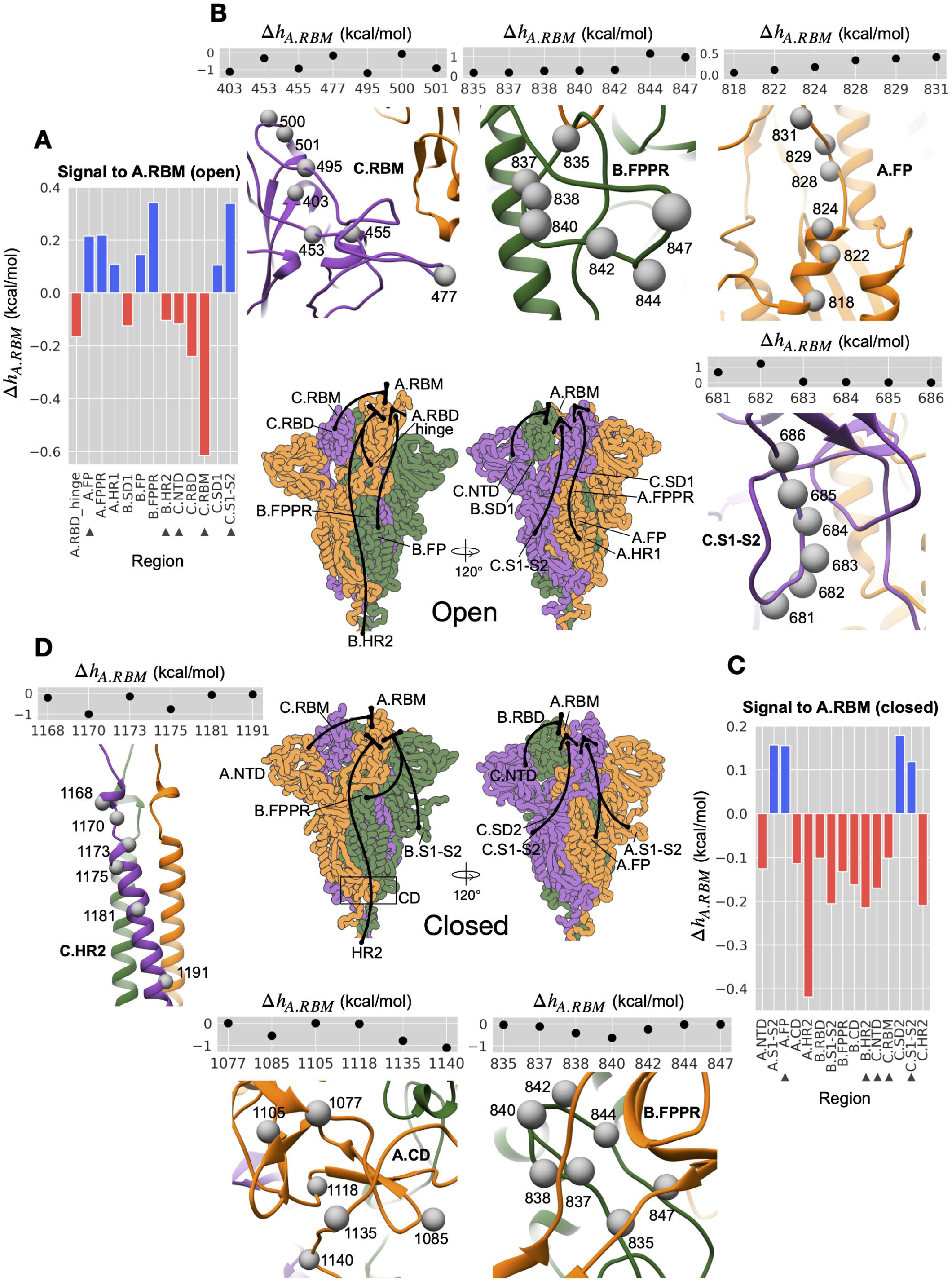
Allosteric signalling to the receptor-binding motif. **(A)** The average modulation ranges at A.RBM residues upon single mutation in different regions (except the A.RBM itself and A.RBD) in the open state. **(B)** Illustration of the effects of signalling from individual residues along with their average modulation ranges in the A.RBM residues in the open state. **(C)** Same as (A) for closed state. **(D)** Same as (B) for closed state. The complete data are available in Figure S3. The triangle symbols mark perturbed regions that cause the same modulation mode in both states. The illustration at the center summarizes the signalling targeting A.RBM from various locations.

Although the overall patterns of signalling appear to be similar between the two states, some distinctions can also be observed. For example, stabilization of B.FPPR, located below A.RBD and A.SD1, lead to a positive modulation in A.RBM in the open state but negative modulation in the closed state. The FPPR loop adopts different conformations — packed below SD1 in the open structure (Figure 3B, top panel, center), but sandwiched between the loops connecting SD1 to SD2 and the helices forming the S2 core in the closed state (Figure 3D, bottom panel, right). Furthermore, a study has pointed out that the disorder-to-ordered transition of FPPR facilitates the distal RBD in adopting the down conformation (Cai et al., 2020), consistent with the observed negative modulation in A.RBM (Figure 3C) and the down A.RBD (Suppl. Figure 2B) upon a stabilizing mutation at B.FPPR in the closed state. On the other hand, positive configurational work exerted in A.RBM (Figure 3A) and the up A.RBD (Figure 2A, 2B) is apparently indicative of conformational changes in the RBD from the up to the down form. Other differences can also be identified: positive modulation by the cluster of A.FP, A.FPPR, A.HR1 and C.SD1 regions in the open state, while only A.FP positively modulates A.RBM in the closed state; the A.RBM is more strongly modulated by HR2 in the S2 stalk in the closed state compared to the open state. Trends in signalling to A.RBM from the same sites of chains B and C are, in general similar, but become weaker upon movement from B to C (Figures 2A and S2).

We next turned our focus to signalling at the single-residue level, and potential effects on the RBMs in the open and closed forms of the S protein. Some of the residues in the RBM-modulating regions are shown as an example (Figure 3B, 2D), along with the mean modulation values at A.RBM upon their mutations (averaged from the modulation range of the A.RBM residues). The complete ASM data is presented in (Figure S2 A, B), and are available for download through the designated page in the AlloMAPS database (Tan et al., 2019) (link will be available upon peer-reviewed publication). In the open state, stabilizing perturbations at residues 403, 455, 495 and 501 in C.RBM, individually, prevent conformational changes of A.RBM (up RBD, Figure 3B). Of note, mutations at residue 501 are known to occur in variants of concern (VOCs). On the other hand, perturbations of residues 828, 829 and 831 in A.FP induce a positive modulation at about 0.5 kcal/mol, while perturbing residues 844 and 847 in B.FPPR and residues 681 and 682 in C.S1-S2 results in even stronger (about two-fold) positive modulation. In contrast, Figure 3D illustrates that residues 838 and 840 in B.FPPR, 1135 and 1140 in A.CD, and, 1170 and 1175 in C.HR2 can elicit strong negative modulation to A.RBM in the closed state upon stabilizing perturbations.

### Glycosylation as a source of allosteric signalling

Viral glycosylation is known to play important roles ranging from protein stability and folding to immune evasion, host tropism and receptor binding (Vigerust and Shepherd, 2007). It is increasingly recognized that the glycosylation of proteins not only impacts protein function orthosterically, but may also give rise to changes in the structure and/or dynamics at distal protein sites via allosteric signalling (Nussinov et al., 2012). The glycosylated residues are indicated on the open spike structure shown in Figure 4A (middle). Although post-translational modifications are not explicitly modelled in the framework of SBSMMA, covalent linkage of a bulky glycan to a residue would generally stabilize the perturbed residue and its vicinity. The latter can be qualitatively mimicked by the effects of amino acid substitution from a small to a bulkier residue, which we use here to estimate the allosteric modulation caused by the glycosylation at a position in the structure. To investigate the potential allosteric effects of the extensive glycosylation (with 22 N-linked glycosylation sites in each monomer (Watanabe et al., 2020a)) in the S glycoprotein, we analyzed the intra- and inter-monomer allosteric signalling from each glycosylated position. The signalling varies between the open (Figure S4A) and closed (Figure S4B) states, as well as between corresponding positions in different monomers in some cases (see Figure S4 for complete data). Several positions mostly located in the lower part of the S2 bulk, such as residues 709, 717, 801, 1074, 1098 and 1134 (Figure S4), cause weak modulation to all protein sites upon glycosylation. On the other hand, residues B.74 and A.234 (Figure 4A, first row) in NTD generally cause positive (arrows) modulation in S1 and negative (bases) modulation in S2 upon glycosylation. The N234 glycan is involved in stabilizing RBD in the up conformation (Casalino et al., 2020), while N74 glycan is located nearby the NTD antigenic supersite potentially affecting the antibody binding (Cerutti et al., 2021; McCallum et al., 2021). It points to a potentially specifically interesting role of N74/234 glycans, because in agreement with a previous work (Cerutti et al., 2021), we did not detect any allosteric signalling from the antibodies binding to the supersite of the N-terminal domain (NTD, (Lok, 2021). Glycosylated residues C.61 and B.61 (middle row) result in a positive modulation in both S2 cores (stronger in closed state), whereas the S1 domains are stabilized. Glycosylation of position 1158 in any of the monomers consistently leads to strong allosteric modulation in the open and closed S glycoprotein (Figure 4B) chiefly similar in all sites (Figure 4A, bottom row) except the HR2 stalk, where negative and positive modulations in the open and closed states were observed, respectively.

**Figure 4.**
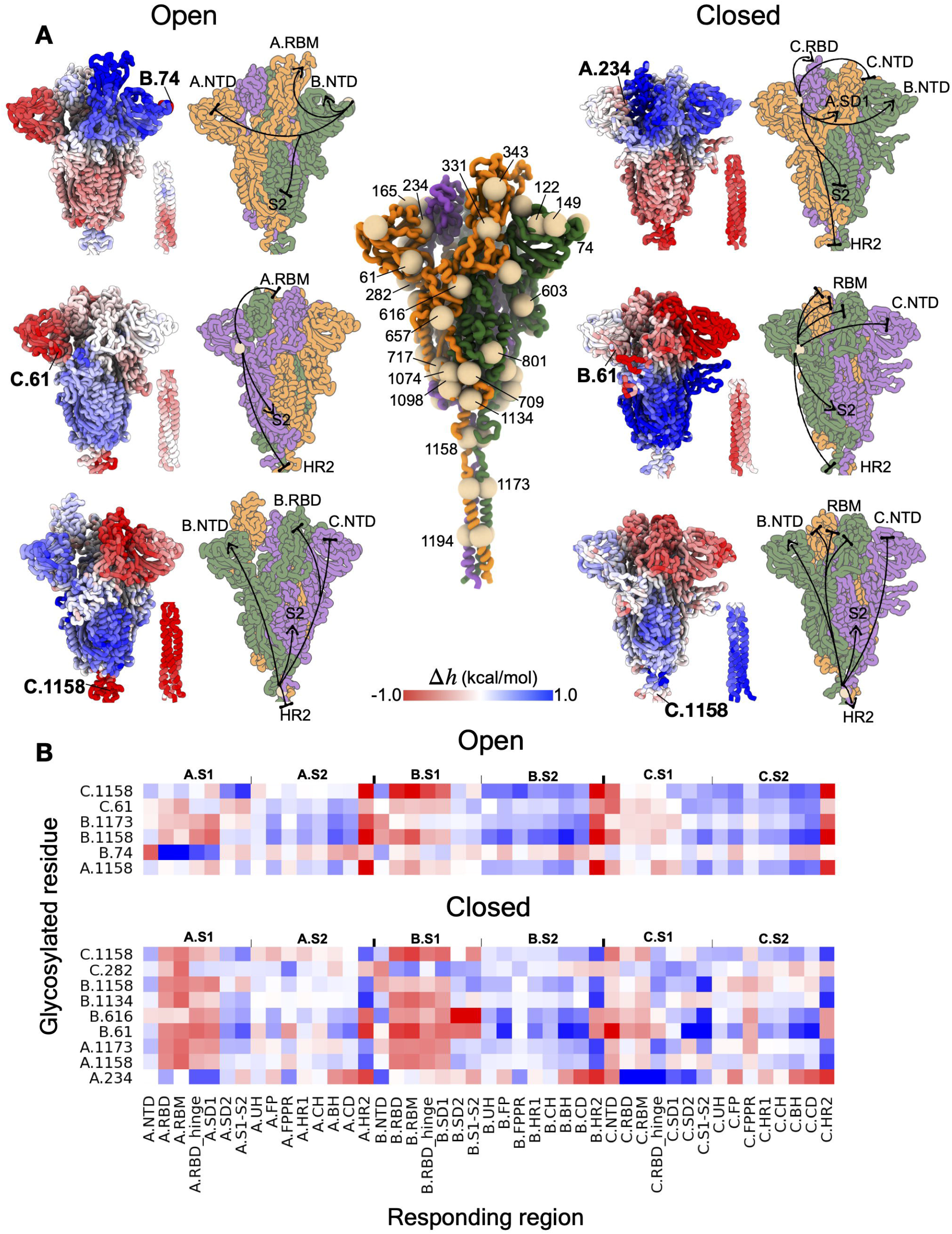
Allosteric signalling from glycosylated positions. **(A)** Center, glycosylated residues are depicted as spheres on the open spike homotrimer. Sides, examples of allosteric modulation originated from different glycosylation sites. The average modulation range 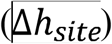 of residues within every responding site/region is provided in full in Figure S4. **(B)** The glycosylated positions that can cause strong modulation (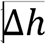, kcal/mol) in multiple sites/regions (with the average 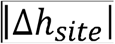 in all sites/regions above 0.2 kcal/mol.

### Allosteric effects of mutations: from an analysis of known mutations to the prediction of potential new variants of concern (VOCs)

The high mutability and rapid evolution characteristic of RNA viruses (Steinhauer and Holland, 1987) prompt a special interest in investigating effects of mutations in the S protein, in general, and those in the RBD, in particular, as the latter is a key factor in the viral entry and in the neutralizing activity of antibodies. The emerged RBD mutations in the widely-circulating VOCs, including Alpha (B.1.1.7, first reported in the UK), Beta (B.1.351, South Africa), Gamma (P.1, Brazil) and Delta (B.1.617.2, India) have been shown to enhance binding affinity to the ACE2 receptor and/or to permit antigenic escape (Gu et al., 2020; Jangra et al., 2021; Rees-Spear et al., 2021; Starr et al., 2020; Wang et al., 2021). Acknowledging that every mutation may have a specific ratio between its ortho- and allosteric modes of action, it remains to be seen if these mutations allosterically affect different S protein regions, in addition to their roles in ACE2 binding and/or immune escape. In general, only 1 out of 6 amino acid substitutions in the spike of the Alpha variant and about half of the substitutions of the Beta, Gamma, and Delta variants occur in the RBD, calling for a comprehensive analysis of all mutations. A striking example is the D614G mutation in SD2, which promotes the distant RBD in adopting open conformations (Benton et al., 2021; Korber et al., 2020; Mansbach et al., 2021). Therefore, it is important to investigate the allosteric effects of recurrent mutations in the whole homotrimer and to uncover latent mutations with a potential to impact the spike’s functions allosterically. To this end, we use the concurrent ASMs (see STAR* Methods for definition), showing effects of genetic mutations present in all chains A-C and evaluating the modulation range exerted at every residue in both open and closed states. Below, we briefly describe the strongest cases of allosteric modulation of the S homotrimer on the basis of the mutational data obtained from (i) strongly-modulating positions identified in the ASMs; (ii) high-frequency mutations from the GISAID database (Elbe and Buckland-Merrett, 2017); (iii) mutations acquired by the VOCs and the amino acid changes with respect to the bat coronavirus RaTG13 (denoted as Bat) - the closest known relative of SARS-CoV-2 with 98 and 90% sequence identity for the ectodomain and the RBD(Wrobel et al., 2020), respectively.

#### Agnostic analysis

We started by employing *agnostic analysis* for identifying residues that can induce the strongest modulation (top 10% in the distributions for positive and negative modulation ranges, see STAR* Methods for details) in the entire ectodomain based on its ASM(Tee et al., 2021). The agnostic analysis reveals residues in multiple sites/regions in the spike that can strongly modulate its dynamics, likely leading, thus, to an extensive allosteric response in the structure when mutated. In the open state, almost all of the strongly-modulating residues are located in the RBD and S2 stalk (Figure S5). Noteworthy, residues 417, 501, and 681 identified by the agnostic analysis are mutated in the Alpha, Beta, Gamma and Delta VOCs, as well as glycosylation sites at residues 74 and 1158, indicating that a mutation/glycosylation at these residues can induce a large allosteric response in the open S homotrimer (Figures 5A, S5). Figure 5A illustrates some of the residues identified in the open state, showing the modulation range exerted at every residue. For instance, simulating amino acid substitutions at residue 985, near residues 986 and 987, which are commonly replaced by prolines (2P-mutation (Kirchdoerfer et al., 2018)) via protein engineering to stabilize the pre-fusion spike conformation, causes large negative modulation ranges in the RBDs (Figure 5A, 5C), and likely prevents conformational changes in the apex of S1. At the same time, the mutation could induce conformational changes in the S2 subunit as indicated by the positive modulation ranges. In another example, stabilizing mutations at residue 1084 in CD produce positive configurational work in S2, while locking the A.RBD in its up conformation (−0.56 kcal/mol) and stabilizing the S2 stalk (Figure 5A, 5D). Some of the positions identified by the agnostic analysis such as 1163 and 1167 (marked by a cross symbol in Figure 5A) are frequently mutated, which is further discussed below. In the closed structure, strongly-modulating residues found in various regions are not implicated in the VOCs known so far (Figure S5 contains complete data from the agnostic analysis of both open/closed forms).

**Figure 5.**
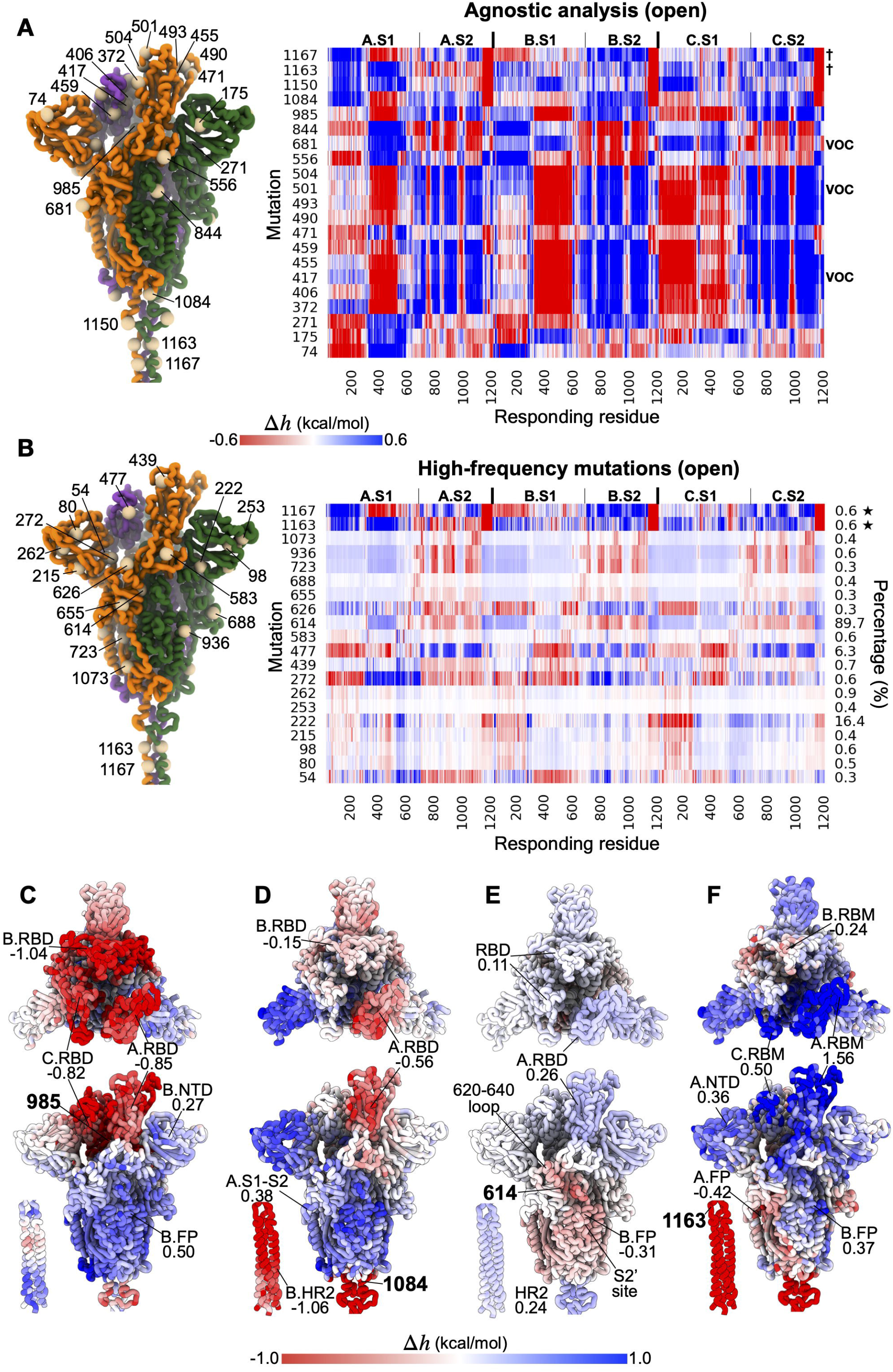
Allosteric modulation detected by the agnostic analysis and caused by high-frequency mutations. **(A)** Left, some of the residues identified from the agnostic analysis on the open spike. Right, the corresponding rows from the ASM are shown (see Figure S5 for complete data). The positions 1163 and 1167 that are mutated in the VOCs are labelled. **(B)** Positions that are frequently substituted by another residue (at least 0.3% out of a total of 218,000 sequences on GISAID, up to 31/11/2020). **(C-F)** The modulation ranges resulted from mutating residues 985, 1084, 614 and 1163 in all monomers with the average modulation range in some of the affected regions indicated.

#### Analysis of the high-frequency GISAID mutations

High frequencies of some S glycoprotein mutations observed in the ongoing pandemic also call for investigating their potential allosteric effects. We identified mutations that occur in at least 0.3% out of a total of 218,000 sequences from GISAID (from the update on 30/11/2020) and showed their modulation values in the open state (Figure 5B). Interestingly, most of these mutations are located throughout the structure and only two frequently-mutated residues (439 and 477) are located in the RBD. Some of these residues cause very weak modulation when mutated including residues 80 and 215 (Beta variant) and 655 (Gamma variant), whereas others such as residues 54, 272, 477, 614, 1163 and 1167 cause relatively strong modulation in multiple regions. Amino acid changes at positions 54 and 272, which are located side-by-side on the β strands near the hinge separating the NTD and other domains in S1, cause positive modulation ranges in the up A.RBD, but negative modulation in the down RBDs in chains B and C (Figure 5B). The S477N mutation at the binding interface with the ACE2 receptor was found to confer resistance to neutralization by multiple monoclonal antibodies (mAbs, (Liu et al., 2021)) and to improve binding affinity to ACE2 (Starr et al., 2020). Our results shows that mutating residue 477 leads to large negative and positive modulation ranges in S1 and S2 subunits, respectively (Figure 5B), suggesting that the escape mutation may also prompt structural and/or dynamical changes in the S homotrimer via allosteric signalling. The dominant form D614G (89.7%) abolishes a salt bridge with K854 of another monomer, and it was found to enhance the infectivity and transmissibility of the virus (Plante et al., 2021; Zhang et al., 2020). A study has shown that the mutation causes ordering of a segment of a disordered loop (residues 620-640 in SD2) which is, in turn, being packed between the NTD and SD1, and stabilizes the furin-cleaved S1-S2 junction (681-686) near the SD2 region that may prevent premature S1 shedding in the D614G mutants (Zhang et al., 2021). The 620-640 loop’s order-disorder dynamics was also correlated with RBD up-down equilibrium and with an effect on the S1-S2 post-cleavage stability (Zhang et al., 2021).We found that simulated stabilizing mutation at residue 614 originates negative configurational work in the 620-640 loop and the S1-S2 cleavage site (Figure 5B, 5E; weaker in A.S1-S2 compared to that in chains B and C). Additionally, negative allosteric modulation was observed in various distal regions forming the S2 fusion core of the metastable pre-fusion structure, including the fusion peptide and the adjacent proteolytic site (S2’ site). At the same time, mutating residue 614 causes a positive modulation in all RBDs, especially in the up A.RBD (Figure 5E). This is consistent with experiments, showing that the G614 trimers predominantly adopt a range of open conformations with at least one RBD pointed upwards, as opposed to the D614 trimer frequently yielding a closed conformation (Benton et al., 2021; Mansbach et al., 2021). The long S2 stalk is also positively modulated, indicating its increased flexibility, which may allow the spike ectodomain to scan the host cell surface more efficiently. We found that the frequently-mutated residues 1163 and 1167 in the S2 stalk, which are identified by the agnostic analysis (indicated by a star symbol in Figure 5B), can strongly modulate various regions of the trimer. For instance, rigidifying the stalk by mutating residue 1163 causes large positive modulation ranges throughout the bulk of the ectodomain (Figure 5F).

#### Variants of concern from the perspective of the evolution from the bat coronavirus RaTG13

Considering the evolution of SARS-CoV-2, we analyzed mutations acquired by the Alpha, Beta, Gamma, and Delta variants of concern and/or substituted in the bat coronavirus RaTG13 (VOCs/Bat), the most interesting of which from the perspective of allosteric signalling are exemplified in Figure 6A (open state; complete data for both open and closed states are available Figure S6A). It appears that some of the mutated residues in VOCs/Bat can induce allosteric responses in the S ectodomain. Stabilizing mutations at residues 417 (Beta/Gamma), 484 (Beta/Gamma/Bat), and 501 (all of VOCs/Bat except Delta) in the RBD cause negative and positive modulations in the S1 and S2 subunits, respectively. On the other hand, a mutation at residue 681 (Alpha/Delta) can cause a large positive configurational work exerted in all RBDs in the open S homotrimer (Figure 6A), whereas a rather mixed response among the RBDs in the closed state can be observed (Figure S6A), suggesting that it may facilitate multiple RBDs to adopt the up conformation and to bind the receptor once a RBD has shifted up. In particular, residues 417, 501 and 681 can originate the strongest modulation (marked by a star symbol, Figure 6A, right) upon their substitution, in a full agreement with the agnostic analysis (Figure S5). Also, the K417N/T, E484K and N501Y mutations have been shown to confer resistance to antibodies (decreased sensitivity, ↓S symbols in Figure 6A, right) and/or enhance binding affinity to the ACE2 receptor in experimental studies (Gu et al., 2020; Jangra et al., 2021; Rees-Spear et al., 2021; Starr et al., 2020; Wang et al., 2021). Figure 6B and C illustrate the per-residue modulation ranges caused by a mutation at residues 484 or 501, respectively. The large positive modulation ranges in S2, which are also observed in the closed state (Figure S6A), indicate conformational changes induced in the distal S2 subunit including the FP, the S1-S2 and S2’ cleavage sites, and the membrane-proximal end of the S2 stalk upon a mutation at residue 484/501, likely impacting the spike’s functions via allosteric signalling. Compared to residue 484, mutating residue 501 causes a more extensive negative modulation range across the RBDs and slightly stronger positive modulation in most parts of the S2 subunit. A double mutation at positions 484 and 501 largely recapitulates the modulation caused by residue 501 alone (Figure 6D). At the time of writing, the Delta variant (B.1.617.2, originated from India), with acquired mutations at residues 478 and 950 in addition to 681 (also present in Alpha) and 452 (present in Epsilon variant originating from California (Motozono et al., 2021)) is driving a global surge of cases. Interestingly, the T478 amino acid in the wild-type Wuhan strain is reverted to lysine, which is the original residue at position 478 in the closely-related bat coronavirus RaTG13, SARS-CoV, and other SARS-related coronaviruses. While the effects of T478K and D950N in the Delta variant are still unknown, the data presented in Figure 6A shows that substitutions of residues 478 (RBD) and 950 (HR1) widely cause allosteric modulation in both the S1 and S2 subunits. Moreover, some positions with amino acid substitutions from RaTG13 (Bat) such as residues 372, 459, 490, 493 and 501 can induce strong allosteric response upon mutation (Figure 6A). Amongst these positions, the F490S mutation is harbored by the Lambda variant, a variant of interest mainly circulating in South America, while the rest have yet to be reported in emerging strains.

**Figure 6.**
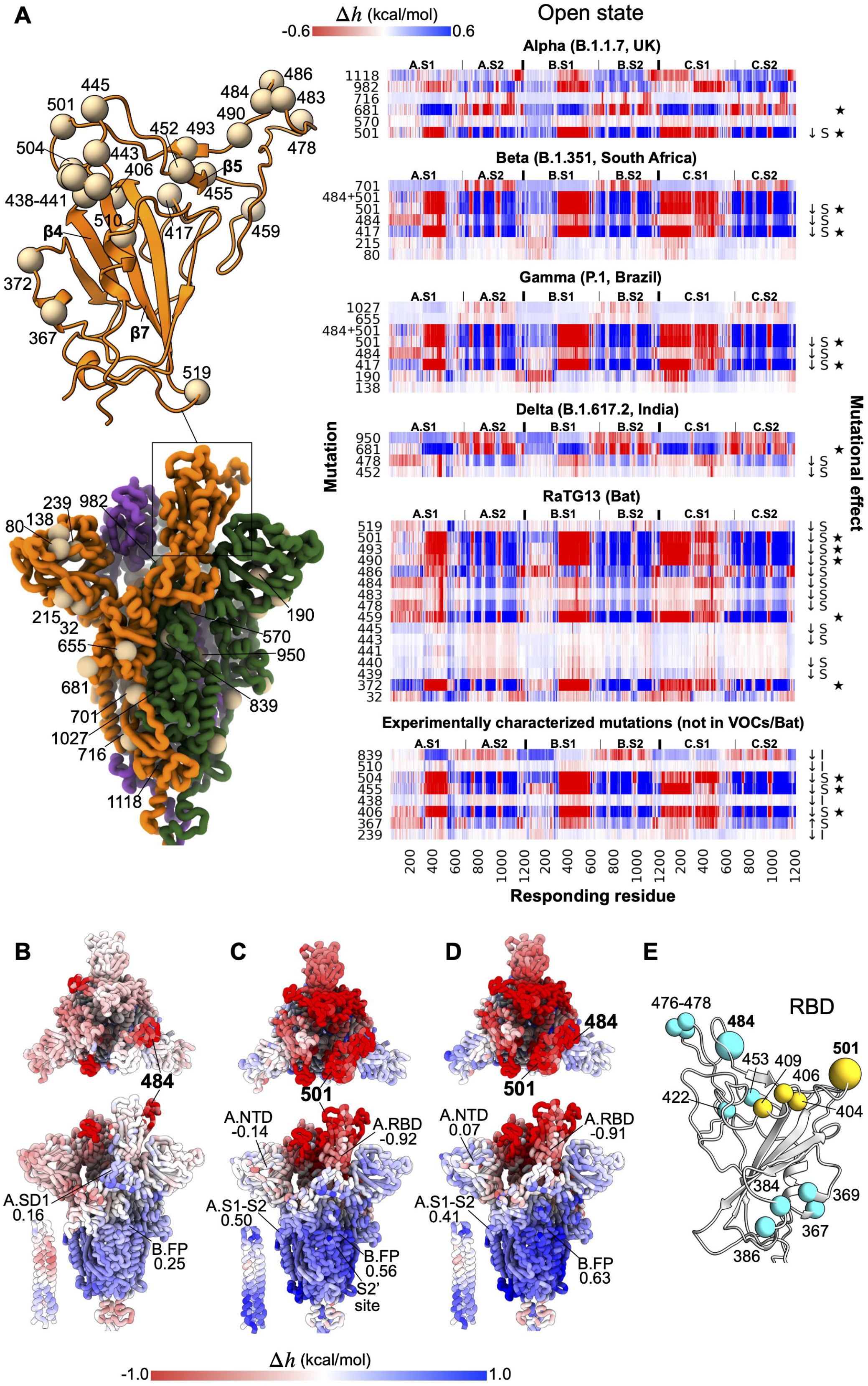
Allosteric effects of mutations from VOCs, Bat, and experiments. **(A)** Right, allosteric modulation ranges of positions that are mutated in VOCs and RaTG13 bat coronavirus. Some experimentally characterized mutations (Li et al., 2020a; Starr et al., 2021) that are not associated with the VOCs/Bat are also included (see Figure S6 for complete data). A mutational effect is denoted by an increase (↑) or decrease (↓) in the infectivity (I) or sensitivity (S) to monoclonal antibodies and/or convalescent sera. Residues identified in the agnostic analysis are marked with a star symbol. Left, the location of the residues in the open spike. **(B-D)** Illustration of modulation ranges caused by mutating residues 484, 501 and both. **(E)** Examples of the allosteric polymorphism for residues 484 and 50. Residues with similar modulation to 484 (cyan) and 501 (yellow) were short-listed if the root-mean-square deviation of per-residue modulation ranges with respect to those from residues 484/501 were below 0.2 kcal/mol, and well separated from residue 484/501 (C_α_-C_α_ distance ≥ 11 Å, see Figure S6B for complete data).

#### From mutations in VOCs/Bat to allosteric polymorphism and new VOCs

Motivated by the obvious allosteric effects of mutations such as 484 and 501, and their apparent involvement in regulation of the S protein function as described above, we tackle here yet another aspect of the allosteric effects of mutations – allosteric polymorphism (Guarnera and Berezovsky, 2020; Tee et al., 2019), *i*.*e*. an existence of a number of positions in the proteins that, upon a perturbation in the form of a certain mutation, produce an allosteric modulation of the function similar to the one originated by already known pathological/regulatory mutations. Using the comprehensive ASM, we exemplify here analysis of the allosteric polymorphism by identifying residues that can exert per-residue modulation ranges comparable to those from positions 484 and 501. Figure 6E illustrates several residues originating similar modulation ranges as residues 484 (colored in cyan) and 501 (yellow); some of them have been shown to affect the viral fitness in experiments when mutated in experiments (see Figure S6B for complete data). For example, spike mutations at position 476 were shown to escape antibodies REGN10933 and LY-CoV016(Starr et al., 2021). Mutations at position 477 are commonly observed (primarily in the 20A.EU2 clade) and the S477N substitution has arisen multiple times independently (Hodcroft et al., 2020), which was reported to enhance ACE2 binding (Starr et al., 2020) and antibody escape (Liu et al., 2021). The Y453F mutation (cyan) has been associated with outbreaks in mink farms and was shown to confer resistance to the REGN10933 antibody (Starr et al., 2021). While mutations at positions 386 and 422 are not yet reported in GISAID (up to 30/11/2020), according to ASM they can, however, exert similarly strong allosteric modulation as position 484 (Figures 6E, S6B). Additionally, with comparable modulation to that by residue 501, the effects of mutations at residues 404 and 406-409 (yellow) are not yet well described, except those of E406W (Starr et al., 2021) and Q409E (Li et al., 2020a), which decrease and increase the sensitivity to antibodies, respectively. Therefore, while the ratio between the ortho- and allosteric components in the effects of mutations should be a topic of future studies, the distal regulatory effects of mutations discussed above coupled with a wide presence of the allosteric polymorphism delineated by the ASM make it rather clear that there might be a number of protein positions that may become major players in newly emerging VOCs.

#### Experimentally characterized mutations

Finally, we considered allosteric modulation caused by mutations in relation to their experimentally characterized effects on viral fitness. To this end, we used data from two high-throughput mutagenesis experiments (Li et al., 2020a; Starr et al., 2021), which altogether identified 60 residues causing changes to the infectivity and/or sensitivity to neutralizing mAbs or convalescent sera. Some of these residues and the modulation ranges are depicted in Figure 6A (bottom right panel), and the complete data for both states are shown in Figure S6C. The ASM revealed that some of the substitutions that allow the RBD to escape recognition by antibodies (decreased sensitivity, ↓S) also induce strong allosteric responses in distal locations in the spike homotrimer. For instance, mutations at residues 455, 504 and 406, each responsible for the decreased sensitivity to antibodies REGN10933, LY-CoV016 and REGN10933 + REGN10987 cocktail (Starr et al., 2021), respectively, cause large positive modulation in the S2 subunit. Moreover, the complete data shows that stabilizing mutations of residues in a RBD-ACE2 interface loop (residues 455-504) generally induce an allosteric response in the spike structure (Figure S6C). In contrast, mutations of residues in the β4-loop-β5 segment and the β7 strand (residues 434-453 and 508-519) of the RBD core, for instance residues 438 and 510, typically result in very weak modulation (Figures 6A, S6C). This suggests an orthosteric mode of action in the case of the S438F and V510L mutations, which lead to decreased infectivity (↓I in Figure 6A, (Li et al., 2020a)).

### Identification of potential allosteric sites

We use the *allosteric probing map* (APM), which is derived by the exhaustive simulation of the binding of a small probe to every consecutive three-residue segment of a protein (Tan et al., 2020), to find potential druggable sites in the spike protein. Specifically, we developed a combined APM/ASM protocol (Tee et al., 2021), including (i) reverse perturbation of allosteric communication for detecting site-to-site coupling; (ii) targeted analysis for estimating the strength of an allosteric signal; and (iii) agnostic analysis for complementing the signalling to a required value (see STAR* Methods for definitions and details).

Here, we sought to uncover potential allosteric sites targeting different spike regions, using the RBM and RBD hinge in S1, FP and the adjacent FPPR in S2, and the HR2 region in the S2 stalk as examples. Figure 7 illustrates a sample of ten potential allosteric sites R1-R10 and the resultant modulation due to ligand binding to them. The APMs for both states are shown in Figure S7, and the complete list of the binding sites along with the modulation values and the solvent-accessible surface area (SASA) are available on AlloMAPS (link and data will be available upon peer-reviewed publication). Ligand binding to the R1 site, between B.RBD and the flanking C.NTD in the closed S structure, causes strong positive modulation in B.RBD, and negative modulation in the A.NTD, B.NTD and the S2 stalk. It was found that biliverdin, a haem metabolite, binds to a cleft on one side of the NTD and inhibits neutralization by antibodies targeting the domain (Rosa et al., 2021). The polysorbate detergent molecule also binds to this cleft (Bangaru et al., 2020). Our result shows that the R2 site, overlapping with the biliverdin binding site, causes similar modulation as the R1 site, suggesting that a recruited haem metabolite in the NTD cleft can induce conformational changes in the down RBD. Noteworthy, the D215G mutation in the Beta variant, P218Q substitution in the RaTG13 bat coronavirus, and the prevalent A222V mutation in B.1.177 and its sub-lineages are all located in the R2 site. All the above hints that R1 site can be considered as a drug target with a potential to overcome an effect of escaping mutations in the biliverdin binding site. The modelled R3 site is formed by residues 617-642 in a disordered loop in the close state. We show that binding to the R3 site, while stabilizing the bound SD2 domain, also causes a large negative allosteric modulation in all RBDs, and results in positive configurational work exerted in the entire S2 subunit. Earlier, it was found that a linear epitope (residue 625-642) in the R3 site may elicit neutralizing antibodies with high specificity (Li et al., 2020b), and potentially druggable hydrophobic pocket underneath the 617-628 loop, overlapping with the R3 site, was also recently discovered (Zuzic et al., 2021), in addition to connection of the 620-640 order-disorder status with the RBD up-down equilibrium and S1-S2 state (Zhang et al., 2021). Simulated binding to the R4 site near the junction between the bulk and the stalk of the S2 subunit likely leads to increased flexibility in the stalk and B.CD. Negative allosteric modulation of the RBD and SD1 of all chains can also be observed. Ligand binding to the R5 and R6 sites at the coiled coil region of the HR2 destabilizes the stalk of the closed spike glycoprotein and the distant NTD of chains A and B, causing negative modulation from the RBD to SD1 in all monomers.

**Figure 7.**
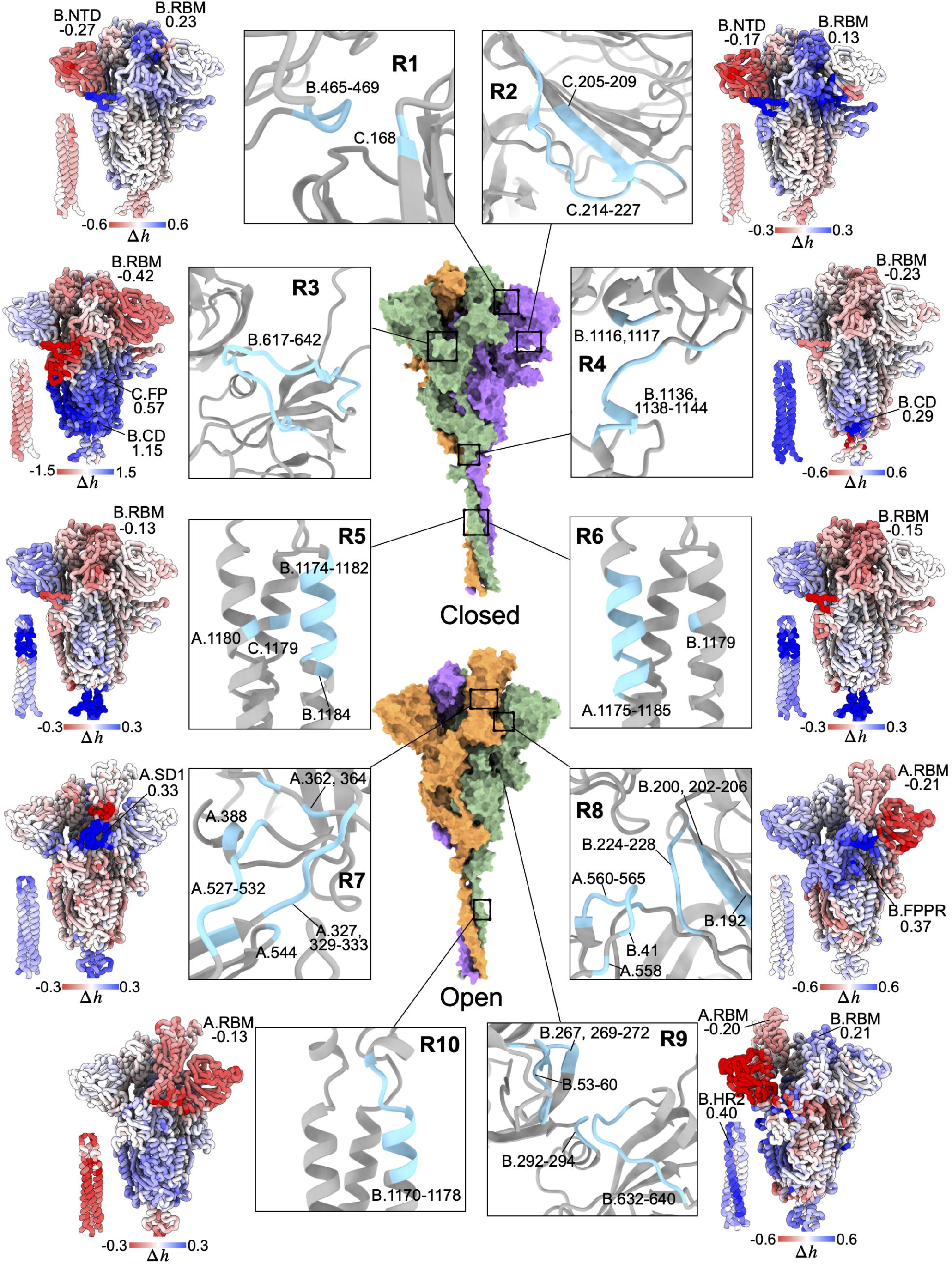
Identification of allosteric sites for prospected drug development. Potential allosteric sites (R1-R10, cyan) identified in the open and closed form of the S protein and effects of signalling (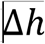, kcal/mol). Allsoteric modulation is in kcal/mol, and it is illustrated by the gradual blue-to-red coloring of structures.The complete list of the identified sites is available in the AlloMAPS database ((Tan et al., 2019), link to downloadable data will be provided upon peer-reviewed publication).

In the open state, binding to the R7 site at the hinge of the up A.RBD chiefly increases the work exerted in the A.SD1, as well as weak negative and positive modulations in the S2 bulk and the stalk, respectively. The R8 site consists of residues from A.SD1 and B.NTD. A linear epitope in A.SD1 (residue 553-564) was discovered to elicit neutralizing antibodies (Li et al., 2020b; Poh et al., 2020). Binding to R8 causes large negative allosteric modulation in the entire B.NTD and slightly weaker modulation in the up RBD, FP and FPPR in chain A. The opposite response can be observed in A.NTD, A.SD1, A.SD2 and B.FPPR. Ligand binding to the R9 site between the NTD and SD2 in chain B negatively modulates the whole B.NTD and the A.RBM, whereas the RBM, FP, FPPR, HR1 and HR2 within the same monomer are positively modulated. Comparing the responses upon ligand binding to the HR2 in the open and closed states showed several differences. First, binding to the R10 site causes negative modulation in the long S2 stalk in the open spike form, whereas R5 and R6 cause the opposite response in the closed state. Second, binding to R10 likely leads to conformational changes in most parts of the S2 subunit, unlike R5 and R6 where mixed responses at S2 are observed.

## Discussion

The current analysis of the S protein’s allosteric regulation spans all levels of its organization and dynamics from considering major structural units of pre/post-fusion states to communication between functional/regulatory sites and down to individual residues. By first, simulating the S protein’s binding to the receptor, we obtained a global picture of the allosteric signalling that facilitates large scale conformational changes of the protein’s subunits (Figure 1). We found that in the open form the conformational dynamics observed in cryo-EM (Cai et al., 2020) and exascale MD simulations (Zimmerman et al., 2021) are supported by the extensive allosteric signalling between the RBMs of different chains, and RBM and other parts, including FP and its S1-S2 cleavage site, and HR2 in the S2 stalk. At the same time, large-scale allosteric communication is strongly suppressed in the closed state of the S protein. Next, on the basis of a per-residue resolution provided by the ASM, we explored details of the allosteric signalling between critical sites of the S protein (Figure 2), observing a strong communication between the RBMs and, specifically, between elements of RBM, RBD and RBD hinges, with FP and HR1/HR2 of the S2 subunit. While in some cases communication between the above sites is persistent in all monomers of both the open/closed states, in others – specific for certain chains in one of the states – we clearly observed an omnipresence and significant signalling between the S1 and S2 parts of the S protein, apparently important for the regulation of its receptor binding and fusion with the cell membrane.

Acknowledging a preventive and therapeutic potential of the drug intervention via a signalling to cause RBM conformational changes incompatible with the receptor binding, we analyzed and described in details perturbations in locations such as RBM (down), S1-S2 cleavage site, FP, FPPR, CD and HR2 that elicit strong allosteric signals to the RBM in the up conformation (Figure 3). Analysis of allosteric signalling from glycosylated residues revealed several positions in both S1 and S2 that may cause a modulation on RBM, NTD, and a few other units important for function or regulation in both the closed and open states (Figure 4).

The high rate of mutations characteristic of RNA viruses and, as a result, their potential contribution to the emergence of new VOCs, requires specific attention to their effects in general and, from the perspective of this work, to its allosteric component. We show here how ASM-based comprehensive analysis combined with available clinical and biophysical data can shed light on the allosteric effects of mutations, allowing to predict potential drivers of new VOCs (Figures 5,6). First, agnostic analysis shows protein positions that originate strong allosteric modulation including residues 417 (Beta/Gamma), 501 (Alpha/Beta/Gamma/Bat) and 681 (Alpha/Delta), and others causing similarly strong modulation as those mutated positions from the VOCs that have emerged, providing a list of potential new candidates (Figures 5A, S5). The predictive power of the SBSMMA is further corroborated by the detection of the allosteric effect of several frequently observed mutations (Figure 5B), such as D614G, which might favor the opening of the spike that leads to increased viral infectivity and transmissibility. While some mutations in VOCs are located in RBD and, presumably, work orthosterically, there are a number of mutations in the RBD acting allosterically and modulating distal parts of the S protein (Figure 6A). In particular, recurrent mutations at RBD residues 417, 484 and 501 in several VOCs could lead to strong allosteric responses in distal functionally important locations in the homotrimer. Notably, strong allosteric signalling provided by residue 501 explains its role in the Alpha, Beta and Gamma VOCs, also pointing to other residues that originate the same signalling being potential “drivers” of new emerging VOCs (*e*.*g*. residues 403, 455, 495; Figure 3B) upon mutation. Analysis of the experimentally characterized RBD mutations reoccurring in different VOCs and RaTG13 bat coronavirus reveals two groups of mutations (Figure 6A): one with allosteric modes of action (417, 478, 484, 486, 490, 493, and 501), and another which works orthosterically (*e*.*g*. 439, 440, 443, 445). We also observed a clear indication of the allosteric polymorphism (Tee et al., 2019), which strongly increases the number of protein positions that could turn into the drivers of new VOCs upon mutation. The complete list of positions that may originate strong allosteric modulation upon mutation is available in the AlloMAPS database ((Tan et al., 2019), link and data will be provided upon peer-reviewed publication).

The existence of allosteric communication between well-separated sites (Figures 1-3) complemented by the allosteric signalling caused by glycosylation (Figure 4) and by the allosteric modulation caused by recurrent mutations (Figures 5,6) point to the potential for utilizing the allosteric mode of regulation for the diagnostic assessment of new VOCs and therapeutic targeting of the S protein. Additionally, increasing evidence of antibodies binding to spike regions outside of the RBD with neutralizing activity accentuates the need to investigate the potential allosteric effects caused by the binding to these locations (Chi et al., 2020; Li et al., 2020b; McCallum et al., 2021) looking towards the design of therapeutic antibodies (Samsudin et al., 2020). Using our APM-based (Tan et al., 2020) generic framework for identifying potential allosteric sites (Tee et al., 2021) on the basis of the reverse perturbation (Tee et al., 2018), we predicted a repertoire of candidate sites for modulating remote functional sites/regions in S glycoprotein (Figure 7). We found several binding sites, which may serve as targets for ligands and neutralizing antibodies (Cao et al., 2020; Chi et al., 2020; Huo et al., 2020; Wrapp et al., 2020; Zhou et al., 2020). Of note, both closed and open states reveal unique potentially druggable sites. For example, in the closed state, the R1 site could be a good drug target candidate, as it originates the same allosteric modulation as the biliverdin-binding site (overlapping with R2), allowing, at the same time, to avoid escaping mutation in the Beta and B.1.177 variants located at the R2 site (Figure 7). The regulatory potential of the R3 site in the closed state is corroborated by the existence of neutralizing antibodies (Li et al., 2020b) elicited by the linear epitope located in it. Decreased dynamics of the RBM, similar to the effect of R3 binding, is observed as a result of the binding to R4-R6, with additional destabilization of the stalk and NTD in the case of binding to R5 or R6. The location of R5 and R6 in the coiled coil region of the HR2 hints to a possibility of using helical stapled peptides (Kim et al., 2011; Kutchukian et al., 2009) as drug candidates for these sites. Remarkably, while the R10 site is found in the open state located in the HR2, similar to R5 and R6 detected in the closed state, binding to R10 causes a strong stabilization of the stalk – the opposite effect to that of binding to R5 and R6. Other examples of binding sites detected in the open state include R7 at the hinge of the up A.RBD, R8 comprised of residues from A.SD1 and B.NTD, and R9 located between the NTD and SD2 in chain B. Binding to these sites causes diverse allosteric modulation in different parts of the S protein, highlighting a potential for drug development. The complete set of binding sites/patches, which may serve as targets for allosteric drugs is available in the AlloMAPS database ((Tan et al., 2019), link and data will be provided upon peer-reviewed publication).

To conclude, while the mutability and the druggability of the S glycoprotein appear to be dichotomous from the perspective of the allosteric regulation of protein functions, both phenomenologies can be viewed as two sides of the same coin, anchored on the cornerstone of protein dynamics (Figure 8). On the “dark side”, the high mutability of viral proteins is a source of a myriad of mutations, some of them with the potential to modulate the structure and dynamics at another location via allostery, leading to phenotypic differences in the variants such as increased infectivity, virulence, or transmissibility (Figure 8). The allosteric mode of action of some mutations makes their analysis challenging, which is further complicated by the wide presence of allosteric polymorphisms and combined effects of several mutations (Tee et al., 2019) that may result in the emergence of multiple new VOCs. On the “bright side”, however, the fundamental property of protein allostery underlined by structural dynamics – remote modulation of protein activity (Guarnera and Berezovsky, 2016a) – can be leveraged to identifying druggable regulatory exosites towards modulation of the sites/regions involved in the protein functions with important therapeutic implications (Figure 8). In the case of the S glycoprotein, it is increasingly recognized that future therapeutic strategies and drug design should target regions beyond the RBD, wherein mutational escape of antibodies have emerged and are spreading rapidly. The lists of mutations and binding patches provided here on the basis of combined ASM-/APM-based analysis of signalling and probe binding can serve as a foundation for future analysis of allosteric signalling and its alteration in the S protein dynamics and for quantification of individual and combined effects of mutations, glycosylation, and binding aimed at the prediction of new VOCs and design of allosterically acting drugs.

**Figure 8.**
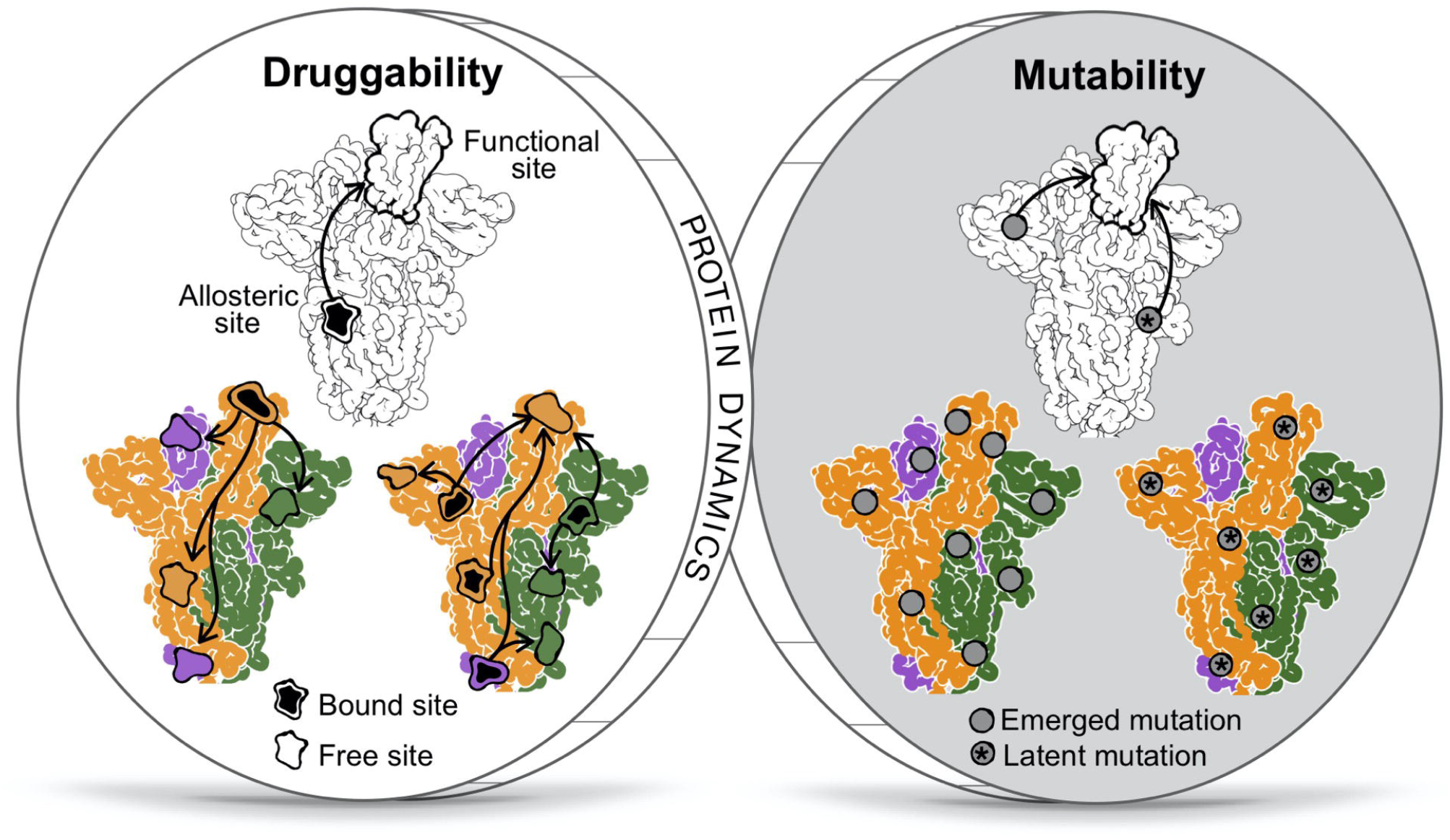
Two sides of the coin: mutability and druggability as a manifestation of the “dark” and “bright” sides of viral protein dynamics. The dark side: Mutation can result in an allosteric response at a site of interest; some of these have emerged and accumulated in circulating VOCs, presumably due to the increased viral fitness, while others are yet to emerge and lead to new VOCs. The bright side: protein dynamics is a cornerstone for the allosteric signalling and modulation of the function from a remote location makes use of the advantages of allosteric drugs.

## Authors contributions

Conceptualization, I.N.B; Methodology, Z.W.T, W.V.T, E.G., and I.N.B; Investigation, Z.W.T, W.V.T, E.G., Z.W.T, F.S., P.J.B, and I.N.B; Software, Z.W.T, W.V.T, and E.G.; Formal Analysis, Z.W.T, W.V.T, F.S., P.J.B, E.G., and I.N.B; Visualization, Z.W.T and W.V.T; Writing – Original Draft, W.V.T, and I.N.B; Writing – Review & Editing, Z.W.T, W.V.T, F.S., P.J.B, E.G., and I.N.B; Supervision, I.N.B.; Project Administration, I.N.B.; Funding Acquisition, P.J.B and I.N.B.

## Declaration of interests

The authors declare no competing financial interests.

## Acknowledgments

Financial support provided by the Biomedical Research Council via Agency for Science, Technology, and Research (A*STAR), is greatly appreciated.

## STAR Methods

### KEY RESOURCE TABLE

#### RESOURCE AVAILABILITY

##### Lead contact

Further information and requests for resources should be directed to and will be fulfilled by the Lead Contact, Igor N. Berezovsky.

##### Materials availability

This study did not generate new materials.

##### Data and code availability

The ASM, APM, and binding sites data have been deposited at the AlloMAPS database and are publicly available upon publication. This paper does not report original code. Any additional information required to reanalyze the data reported in this paper is available from the lead contact upon request.

## EXPERIMENTAL MODEL AND SUBJECT DETAILS

### Building S protein model

Integrative homology modelling was performed to build complete structural models of the S protein in open and closed states using a combination of cryo-EM and NMR structures as templates (details below).

### Collection of characterized mutations

A total of 60 residues known to cause phenotypic changes upon non-synonymous substitutions were obtained from the experimental results by Li *et al*. (Li et al., 2020a) and Starr *et al*. (Starr et al., 2021) to investigate their potential allosteric modulation. For the latter study, we used only those amino acid positions with a total escape of at least 1.0, which indicates antigenic escape from antibodies.

## METHOD DETAILS

### Integrative Homology Modelling

Modeller version 9.21 (Sali and Blundell, 1993) was used to build two full length models of SARS-CoV-2 S protein: i) open state with one RBD in the up conformation and two RBDs in the down conformation; and ii) closed state with all three RBDs in the down conformation. For the open state model, the cryo-EM structure of S protein ECD in the open state (PDB: 6VSB, (Wrapp et al., 2020)) was used as the main template, with missing loops in the NTD and C-terminus of the ECD modelled using the higher resolution cryo-EM structure of S protein ECD in the closed state (PDB: 6XR8, (Cai et al., 2020)). The latter was also used as the main template for the closed state model. The HR2 domain of the S protein stalk was modelled using the NMR structure of SARS-CoV HR2 domain in the prefusion state (PDB: 2FXP, (Hakansson-McReynolds et al., 2006)), which shares 96% sequence identity. The S protein transmembrane (TM) domain was modelled using the NMR structure of HIV-1 gp41 TM domain (PDB: 5JYN, (Dev et al., 2016)), which shares 27% sequence identity. For each state, ten models were built and the three models with the lowest discreet optimized protein energy score (Eramian et al., 2006) were selected for further stereochemical assessment using Ramachandran analysis (Ramachandran et al., 1963). Finally, the best model was chosen as the one with the lowest number of outlier residues.

### Statistical mechanical model of allostery

The structure-based statistical mechanical model of allostery (SBSMMA) introduced and described in previous work (REFs) is used here to quantify the energetics of the allosteric communication in the spike glycoprotein at the single-residue level. Briefly, the effect of ligands, mutations, their combinations, and probes (in every case it is a perturbation *P* of the original state 0) is evaluated at the single-residue resolution as a free energy difference 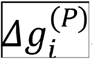. A single crystal structure is used to construct the Cα harmonic model for the unperturbed and perturbed states of the protein. The energy function associated with this perturbed *P* state is 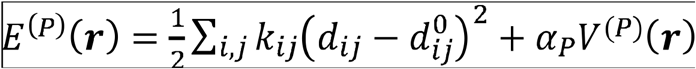. The first term, 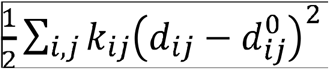, is the energy function associated with the unperturbed state, where 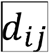 and 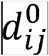 are the interatomic distances between 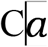 atoms in the generic 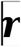 and reference 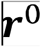 structures, respectively, and 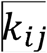 are spring constants. The second term, 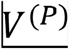, is an additional harmonic term, and where 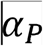 is a perturbation parameter (Tee et al., 2021). Ligand binding perturbations are defined via the over-stabilization of the interactions between all pairs of residues in the binding site, ligand-probe perturbations are accounted via strengthening the interactions between residues of three-residue protein chain segment and nine closest residues with the shortest average distance to these three bound by the probe. Two types of mutations, stabilizing (UP) and destabilizing (DOWN), are considered (Tee et al., 2021). The set of orthonormal modes 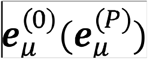, characterizing the configurational ensemble of the original (perturbed) protein state, is calculated from the Hessian matrix 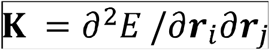. The *allosteric potential* measuring the effect of a perturbation on a particular residue *i* is evaluated as the elastic work exerted on residue *i* as a result of the changes in the neighbourhood conformational ensembles caused by the low frequency normal modes 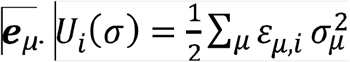, where 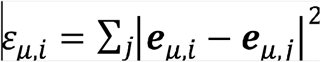 and is 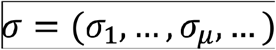 a set of Gaussian distributed amplitudes with variance 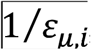, which identifies the displacement of the residue 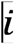 as 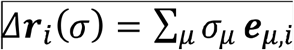. Integration over all possible displacements of neighboring residues provides the per-residue partition function 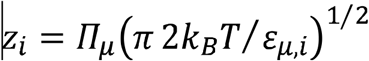 and, then, the *free energy* 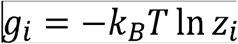 associated with the transmitted allosteric signal. The free energy difference between two (unperturbed/perturbed) protein states quantifies the allosteric signaling delivered to a residue as a result of a perturbation 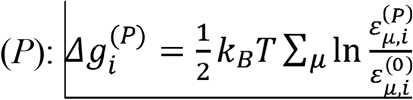. The work exerted due to purely allosteric effect is defined as the *allosteric modulation* 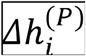, i.e. the deviation of the free energy difference 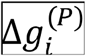 from its mean value over all the residues of the protein (protein chain), containing the residue *i* : 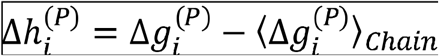. Positive modulation indicates potential conformational changes of residue *i* whereas a negative modulation may prevent conformational changes. It should be noted that the magnitude of the modulation should be considered in relation to the thermal fluctuations. While per-residue modulations with the magnitude above *k*_B_*T* should be regarded as a strong manifestation of allosteric communication, combinations of low-value modulations in a homogeneously affected region may also result in a significant allosteric modulation. The allosteric modulation on the site can be evaluated by the averaging of per-residue modulations 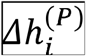 over the residues belonging to this site: 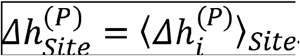.

The *allosteric modulation range* is also used for estimating the allosteric signalling from any position *m* regardless of the original residue in the structure to every residue in the protein and for building allosteric signalling maps (ASMs). It is calculated as a strength of the allosteric signal caused by the substitution from the smallest (Ala/Gly-like) to the bulkiest residues (for example, Phe or Trp), is used as a generic estimate of the allosteric signaling from any protein position *m* to position *i*: 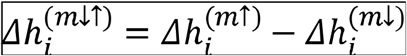. This provides a generic estimate of signaling strength from one position to another, regardless of the original residue at the perturbed position. Depending on the task, two types of the ASMs, concurrent and sequential, can be calculated. In the concurrent ASMs the amino acid changes are simulated simultaneously in corresponding positions in all monomers of the oligomer, allowing to consider effects of genetic mutations. The sequential ASMs, in which amino acid changes are simulated in every monomer one after another, are used as a technique for the analysis of the intra- and inter-monomer signaling from a perturbed position in a certain location of the protein oligomeric structure.

The Allosteric Probing Map (APM), in which the allosteric modulation on residue 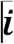 is originated by the binding of a small probe to a three-residue segment of the protein modelled sequentially from residue 1 to residue N-2 for a protein chain of N residues are also derived. The allosteric modulation caused by the probe is also evaluated as a background free effect: 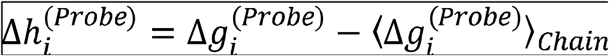.

### Agnostic and targeted analyses of ASM/APM

The agnostic analysis (Tee et al., 2021)of the ASM/APM was carried out to identify 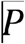 perturbations (mutations and probe binding) that can elicit the strongest positive or negative modulation on distal residues 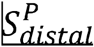 (with C_α_-C_α_ distance above 11 Å) across the protein. Characterizing each perturbation 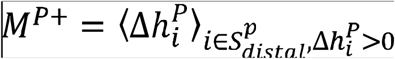 by the average modulation on positively modulated distal residues and negatively modulated residues 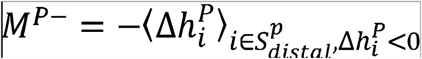, we shortlisted perturbations that ranked in the top 10% for 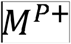 or 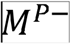. The targeted analysis (Tee et al., 2021) provides a more focused approach, by detecting perturbations that strongly modulate known regions of interest. The modulation of a site/region due to a general perturbation 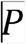 is defined as the mean of the modulation ranges exerted on all residues 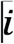 belonging to the site,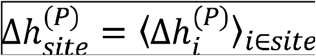. For each targeted site, we ranked perturbations by the allosteric modulation on the given site, and shortlist perturbations in the positive and negative 10% tails for the distribution of 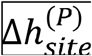.

### Reverse perturbation of allosteric communication

We demonstrated previously that simulating ligand binding to a functional site causes large allosteric responses at the residues in the known allosteric site(s), thereby allowing their identification (Tee et al., 2018; Tee et al., 2021). According to the operational definition of allostery (Tee et al., 2018) in the framework of SBMMA, only a set of distal residues 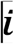 that are beyond a Cα-Cα cutoff distance 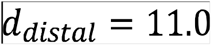 Å away from the site, denoted as 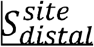, are considered. For each targeted site, we identified distal residues that are strongly modulated by binding on the site, with a modulation 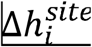 greater than 1.25 times the average magnitude of modulation on distal residues.

### Identification of potential allosteric sites

We have previously introduced a generic framework for the inducing and tuning of allosteric signaling via the identification and design of allosteric sites (Tee et al., 2021). The framework comprises independent components including the agnostic and targeted analyses, and the reverse perturbation of allosteric communication. We obtained a list of potential allosteric sites targeting different regions by combining proximal residues (C_α_-C_α_ cutoff distance of 7 Å) shortlisted in all three approaches. The complete data including the binding sites, the resulting modulation and the solvent-accessible surface area calculated by the FreeSASA package (Mitternacht, 2016) are available on AlloMAPS ((Tan et al., 2019), link and data will be available upon peer-reviewed publication).

## QUANTIFICATION AND STATISTICAL ANALYSIS

No statistical analysis was performed

## ADDITIONAL RESOURCES

No additional resources were used

## Notes

### Competing Interest Statement

The authors have declared no competing interest.

