## Supplementary Figures 1-7 for "Allosteric perspective on the mutability and druggability of the SARS-CoV-2 Spike protein"

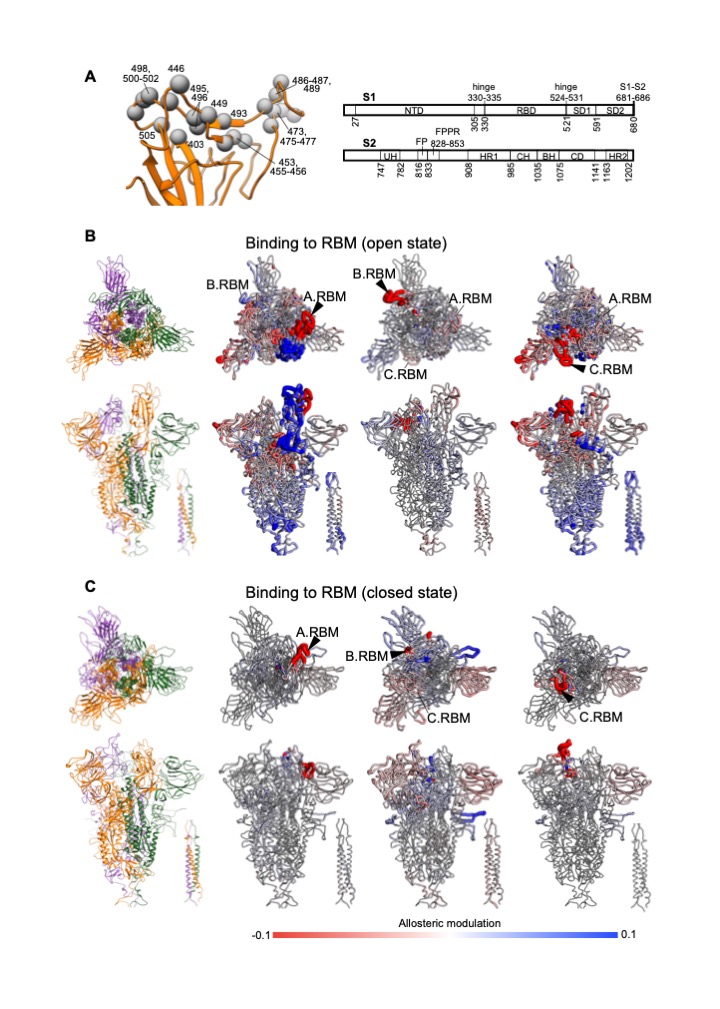

**Figure S1. Definition of the receptor-binding motif (RBM) and the allosteric response upon binding to the RBM.** (A) Left, residues forming the receptor-binding motif (RBM) are shown as spheres on A.RBD of the open spike. The residues are defined based on a C_α_-C_α_ distance cutoff of 5 Å from any residue of the ACE2 receptor, using a structure (PDB: 6M17) of the RBD-ACE2 complex. Right, the residues forming each studied spike region. (B, C) Allosteric modulation of the spike in the open and closed states upon simulated binding to a RBM (marked by a triangle).

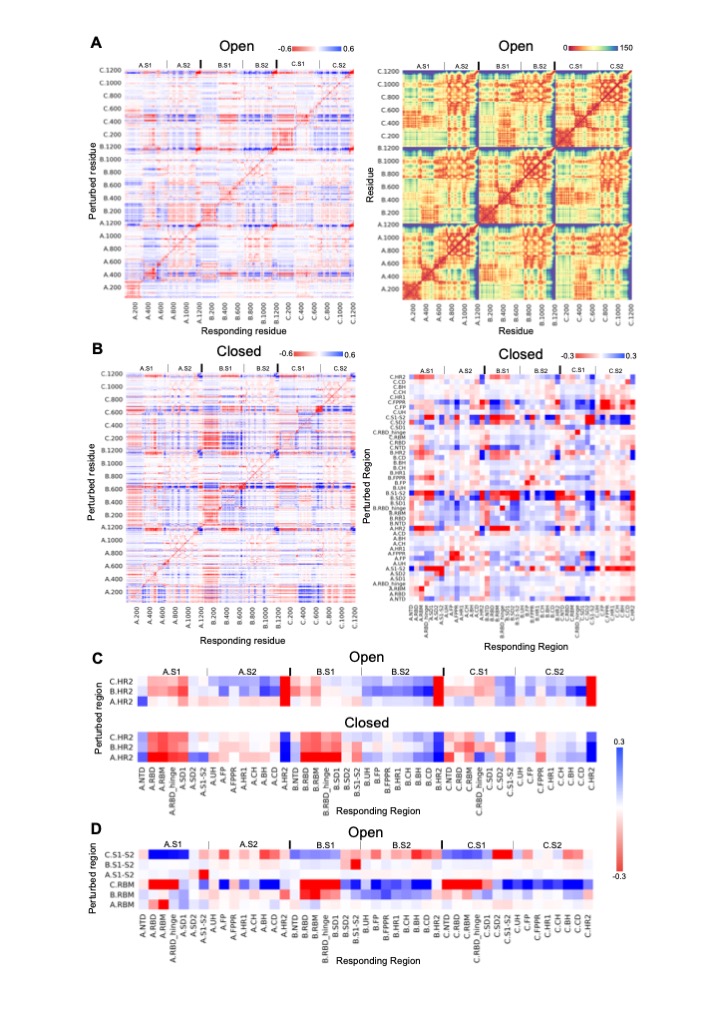

**Figure S2. The complete allosteric signalling maps and some relevant parts.** (A) ASM (left) and the pairwise C_α_-C_α_ distance map (right, Å) of the open spike. (B) The ASM for the closed spike at the per-residue and per-site levels. (C, D) The parts of the ASMs (Figures 2A and S2B, right) corresponding to the results described in Figure 2D and 2E.

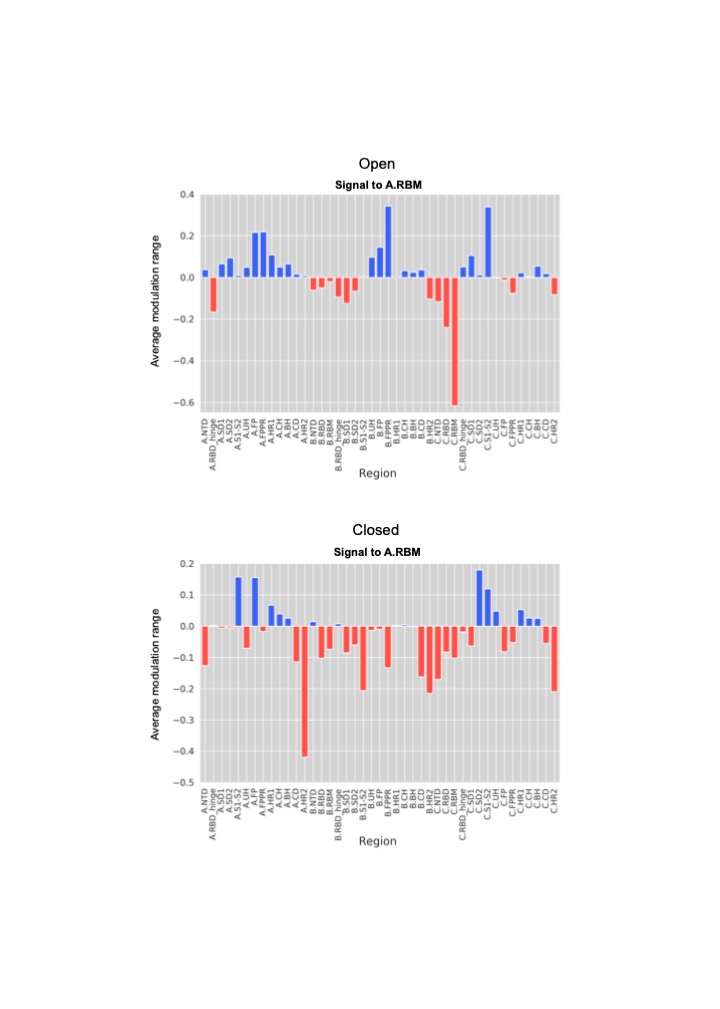

**Figure S3. The average modulation range in A.RBM caused by single mutations in every region (except A.RBM and A.RBD).**

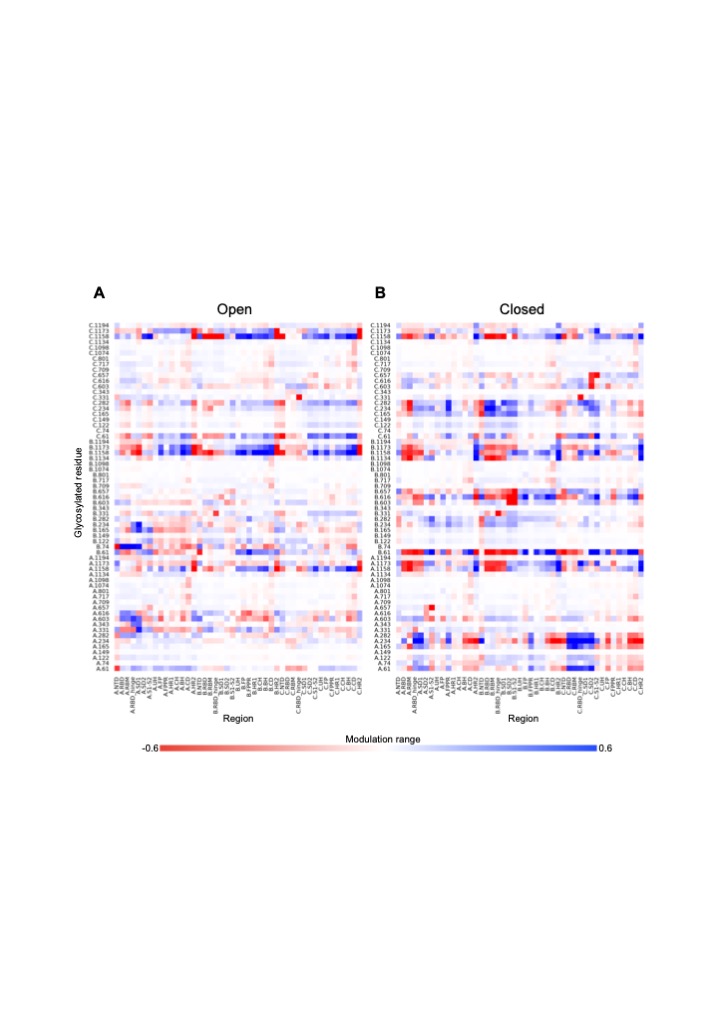

**Figure S4. The average modulation range that resulted in every responding site/region due to signalling from each glycosylated position in the open/closed S homotrimer.**

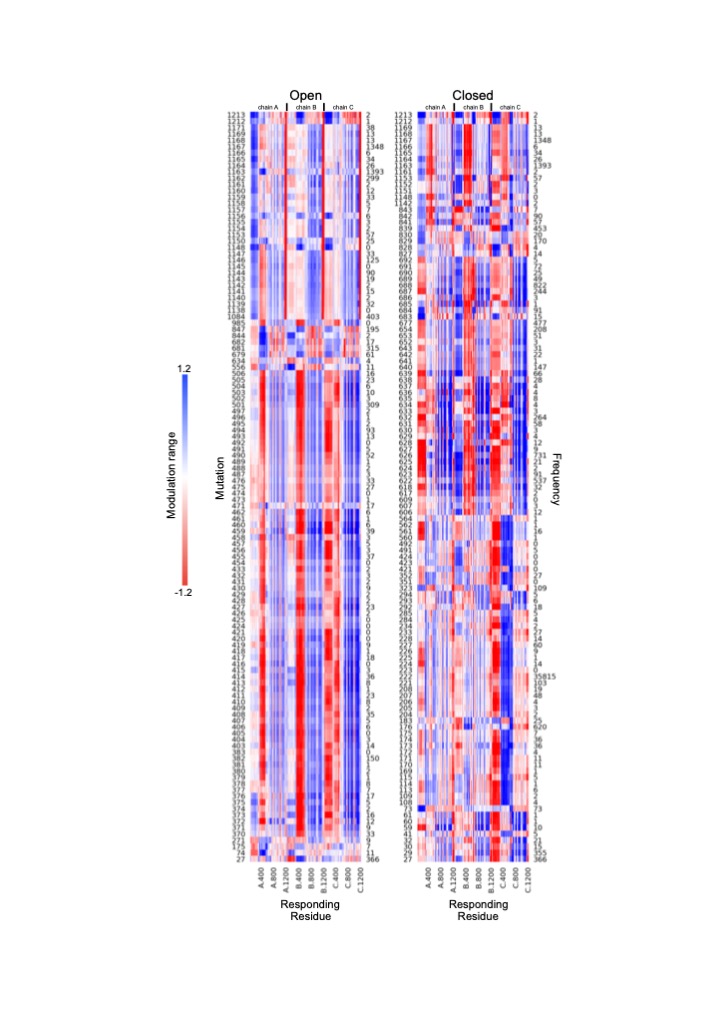

**Figure S5. Agnostic analysis of the ASMs.**

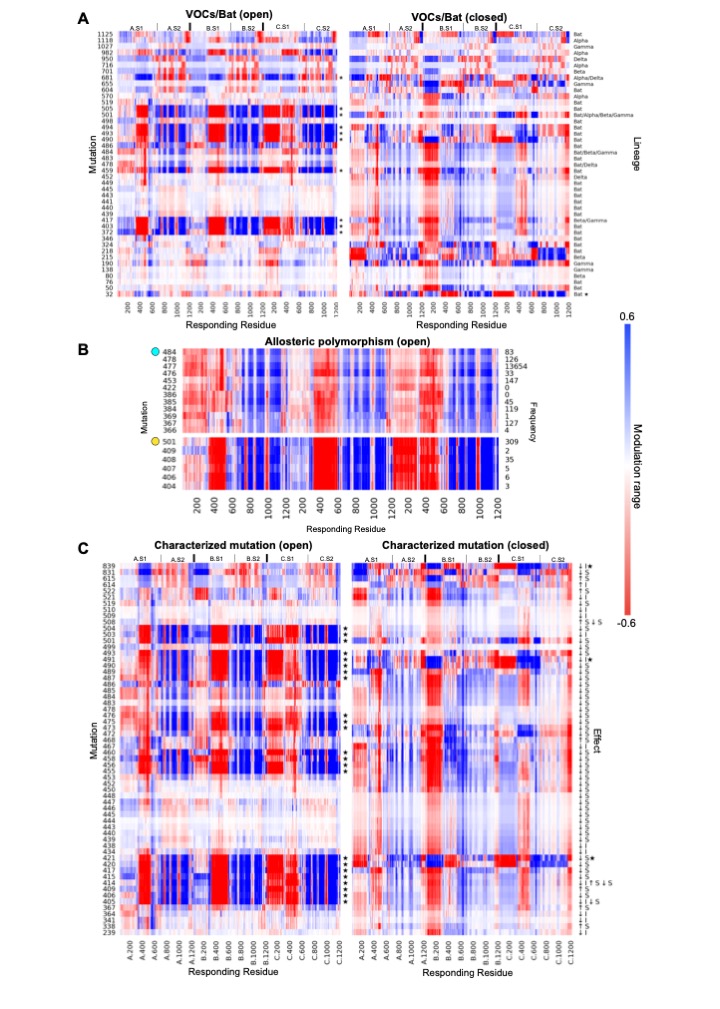

**Figure S6. Modulation ranges caused by mutations in open/closed spike.**

Modulation ranges originated from (A) mutated positions in VOCs/Bat, (B) positions 484/501 as well as those causing a similar response, and (C) experimentally characterized mutations.

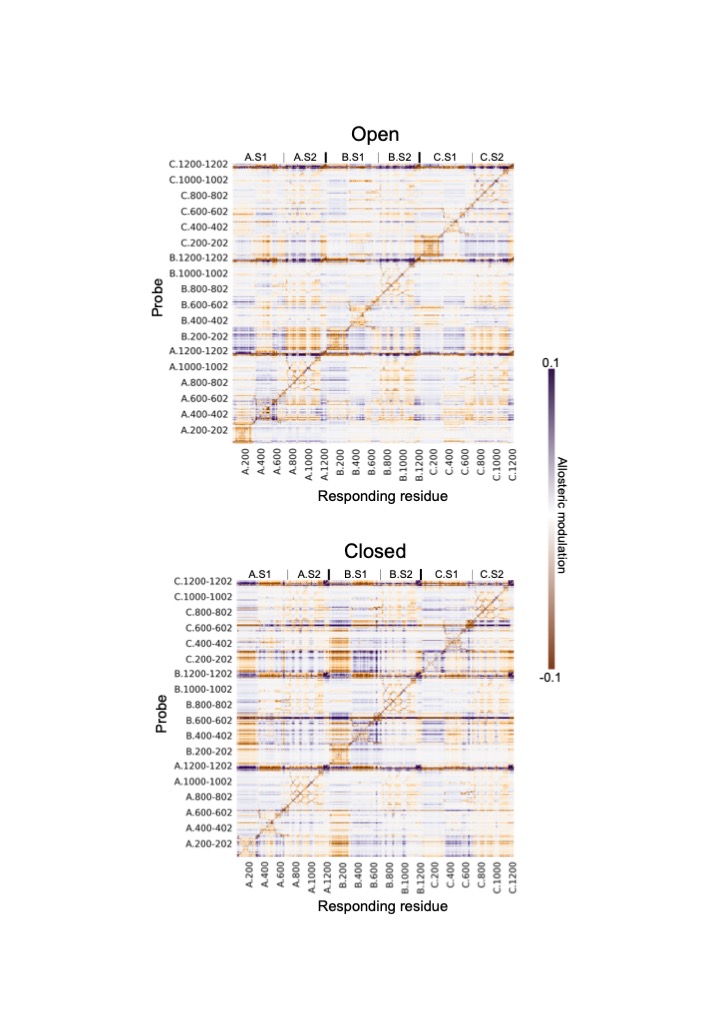

**Figure S7. Allosteric probing maps.**
